# Base editing of *EIF4E* creates novel resistance alleles against bymoviruses in winter barley

**DOI:** 10.64898/2026.09.04.749431

**Authors:** Robert Eric Hoffie, Antje Habekuss, Dragan Perovic, Iris Hoffie, Frank Ordon, Jochen Kumlehn

## Abstract

The barley yellow mosaic virus disease is one of the most important threats of barley production in Europe and Asia. Transmitted by the soil-borne plasmodiophorid *Polymyxa graminis*, there are no direct control options against the causal bymoviruses *Barley Yellow Mosaic Virus* (BaYMV) and *Barley Mild Mosaic Virus* (BaMMV). Resistance breeding is thus the only viable approach and has been very successful in the past, with the resistance-conferring alleles *rym4* and *rym5* of the EUKARYOTIC TRANSLATION INITIATION FACTOR 4E being used extensively in European winter barley breeding. However, virus strains have meanwhile overcome this resistance. Therefore, there is an urgent need for new sources of resistance. Genome editing with Cas endonucleases is a timely and promising approach in this respect. However, the small insertions and deletions that frequently arise during site-directed mutagenesis usually lead to the knockout of the target genes. In the case of *EIF4E*, loss-of-function is accompanied by significant yield reduction. Consequently, more precisely edited alleles with retained function are necessary for crop improvement. The present study represents the first application of base editing in barley plants, using the *EIF4E* gene as an example. Base exchanges were made at two positions in this gene using an nCas9-cytidine deaminase fusion, resulting in a total of 10 novel *EIF4E* alleles in addition to the introduction of a single nucleotide polymorphism that is part of *rym4*. Two of these newly generated alleles led to resistance upon BaMMV inoculation without adverse effects on yield, proving this approach promising to generate new material for resistance breeding.

## Introduction

The EUKARYOTIC TRANSLATION INITIATION FACTOR 4E (*EIF4E*) is a well-known susceptibility factor for infection with bymoviruses in a broad variety of plant species (reviewed in Singhal et al., 2026). In host cells, eIF4E binds to the m7G-cap of eukaryotic mRNA to facilitate the formation of the translation initiation complex, forming the translation loop and let the smaller subunit of the ribosome bind to the mRNA as the starting point for translation of the mRNA’s information into the amino acid sequence (Mars et al., 2024). Bymoviruses belong to the family of Potyviridae that are single stranded (ss)+RNA viruses infecting plants through transmission by vectors such as aphids or fungi. In host plant’s cells, the virus unpacks its RNA genome and mimics the m7G cap structure with its viral genome-linked protein (VpG) at the 5’ end of its RNA. By that, the viral RNA is recognised by eIF4E as endogenous plant mRNA, whereupon translation of the viral proteins is initiated. The virus then multiplies and spreads throughout the plant (Singhal et al., 2026).

Barley was one of the first plant species where *EIF4E* was identified as a bymovirus susceptibility factor and resistance-conferring alleles were described (Stein et al., 2005; Kanyuka et al., 2005). A screen of several hundred barley accessions from gene banks revealed that all spontaneously evolved resistance alleles carry non-synonymous single nucleotide polymorphisms (SNPs) in their coding regions, leading to an altered amino acid sequence in the binding domain of eIF4E, which disturbs the interaction with the viral VpG RNA cap (Yang et al., 2016). Two of those alleles, namely *rym4* and *rym5*, are widely used in European winter barley breeding and most of the resistant cultivars rely on that resistance mechanism. Due to high selection pressure, first strains of *Barley Mild Mosaic Virus* (BaMMV) and *Barley Yellow Mosaic Virus* (BaYMV), the economically most important bymoviruses in barley, have overcome the *rym4*/ *rym5*-mediated resistance, which is why barley breeding is in need of new sources for resistant cultivars.

For several plant species, the targeted knockout of *EIF4E* or its isoform EIF(iso)4E was reported to generate bymovirus-resistant plants (reviewed in Schmitt-Keichinger (2019)). By targeted mutagenesis of *EIF4E*, knock-out lines were generated in barley and proven resistant to mechanical inoculation with BaMMV. However, these lines were impaired in yield parameters, suggesting that the functions of *EIF4E* for the plant cannot be fully replaced by its isoform *EIF(iso)4E* (Hoffie et al., 2021b). On the other hand, in regards to resistance-breaking virus strains, new alleles of *EIF4E* are desirable. To generate a variety of new, functional alleles of *EIF4E* with single nucleotide polymorphisms (SNPs), more precise mutagenesis approaches appear promising.

Inducing mutations by means of CRISPR-associated endonucleases at specifically targeted sites in the genome, resulting in largely random modifications of the nucleotide sequence, is the state-of-the-art approach to application-oriented genome editing in the vast majority of crop species including barley (Hisano et al., 2021). However, base editing is an emerging tool that allows for more precise genetic alterations.

Base editing was first described with the combination of a cytidine deaminase with the Cas9 endonuclease by Komor et al. (2016). These authors tested different combinations of cytidine deaminases (human AID, human APOBEC3G, rat APOBEC1 and sea lamprey CDA1) and dCas9 (with deactivated endonuclease domains) as well as nCas9 (a nickase variant of Cas9 with deactivated RuvC nuclease domain by D10A mutation) in human cell lines. Shortly afterwards, Nishida et al. (2016) presented an alternative system used in yeast (*Saccharomyces cerevisiae*). They fused an nCas9 with an activation-induced cytidine deaminase (AID) orthologue, CYTIDINE DEAMINASE 1 (CDA1) from sea lamprey (*Petromyzon marinus*). Cytidine deaminases are involved in the immune system of vertebrates on the one hand in rendering viral DNA harmless by mutation, but on the other hand, they also act in hypermutation in B cells in the adaptation and maturation of antibodies (Knisbacher et al., 2016). CDA1-nCas9 fusions are directed to the genomic target motif by gRNA complementarity, while the helicase activity of nCas9 makes the target site accessible for editing. Notably, CDA1 can only modify single stranded DNA. The cytidines within an area of about 5 bp at the 5’ end of the target motif (the editing window) can be subjected to editing.

The processed cytidines are first converted into uridines by deamination, resulting in U:G mismatches between the edited and the complementary strands. The removal of the induced uracil by the uracil DNA glycosylase and the associated base excision repair is prevented by an uracil DNA glycosylase inhibitor (UGI) from *Bacillus subtilis* that is fused to the CDA1-nCas9 chimeric enzyme in addition. Triggered by the single-strand break of the non-edited strand by nCas9, the mismatch repair is activated, which replaces the guanine with adenine on the complementary strand. During the following DNA replication, uridines are misinterpreted by the DNA polymerase as thymines and T-A base pairs are incorporated into the DNA double strand accordingly. Thus, C-to-T (or G-to-A on the complementary strand) substitutions can be induced (Nambiar et al., 2022). These Cas9 derivatives, sometimes referred to as cytidine base editors (CBEs), have been successfully used in various di- and monocotyledonous plants, including Arabidopsis, canola, tomato, rice and wheat (Chen et al., 2017; Hu et al., 2020; Kashojiya et al., 2022; Li et al., 2017; Zong et al., 2017, comprehensively reviewed in Li et al., 2023).

Shortly after the emergence of cytidine base editors, adenine base editors (ABEs), i.e. fusions of nCas9 and an adenine deaminase, were also reported. These are capable of inducing A-to-G substitutions, which was first demonstrated in human cells (Gaudelli et al., 2017). Unlike cytidine deaminases, there are no known DNA-modifying adenine deaminases that have emerged through evolution. Therefore, Gaudelli et al. (2017) developed the DNA adenine deaminase ecTadA* by targeted evolution from the *E. coli* RNA adenine deaminase (ecTadA) that is active *in vivo* on tRNAs. Fused to an nCas9, ecTada-ecTadA* dimers deaminate adenosines to inosines in an editing window of about 4-6 nucleotides around positions 4 to 7 of the target motif. Through complementary base pairing with cytidine, inosine is read and implemented by DNA polymerases as guanine during DNA replication (Nambiar et al., 2022). In plants, adenine deaminases have been used so far in rice, wheat, and in *Nicotiana benthamiana* (Hua et al., 2018; Li et al., 2018; Wang et al., 2021b, comprehensively reviewed in Li et al., 2023).

Here, we present a base editing approach employing CDA as well as ADA base editors to induce targeted SNPs in the *EIF4E* gene to generate new alleles. In total, three and two resistance-associated positions were targeted with CDA and ADA, respectively. Inheritance of induced base edits was followed until M_4_ generation and mutant plants were inoculated with BaMMV to test for resistance. To identify those alleles that combine virus resistance with high plant performance, yield components were assessed.

## Material and Methods

### Selection of target motifs

To induce amino acid substitutions in the eIF4E binding domain, target motifs within the barley *HvEIF4E* gene (HORVU.MOREX.r3.3HG0327270) on chromosome 3H were selected manually in four regions where base substitutions had previously been identified in the resistance alleles *rym4*, *rym5*, *HOR3298 and HOR4224* (Shi et al., 2019; Perovic et al., 2014; Stein et al., 2005). Within these regions, a total of five target motifs were identified and addressed either with cytidine deaminase (CDA) or adenine deaminase (ADA), see Figure 1 and sequences S1.

**Figure 1:**
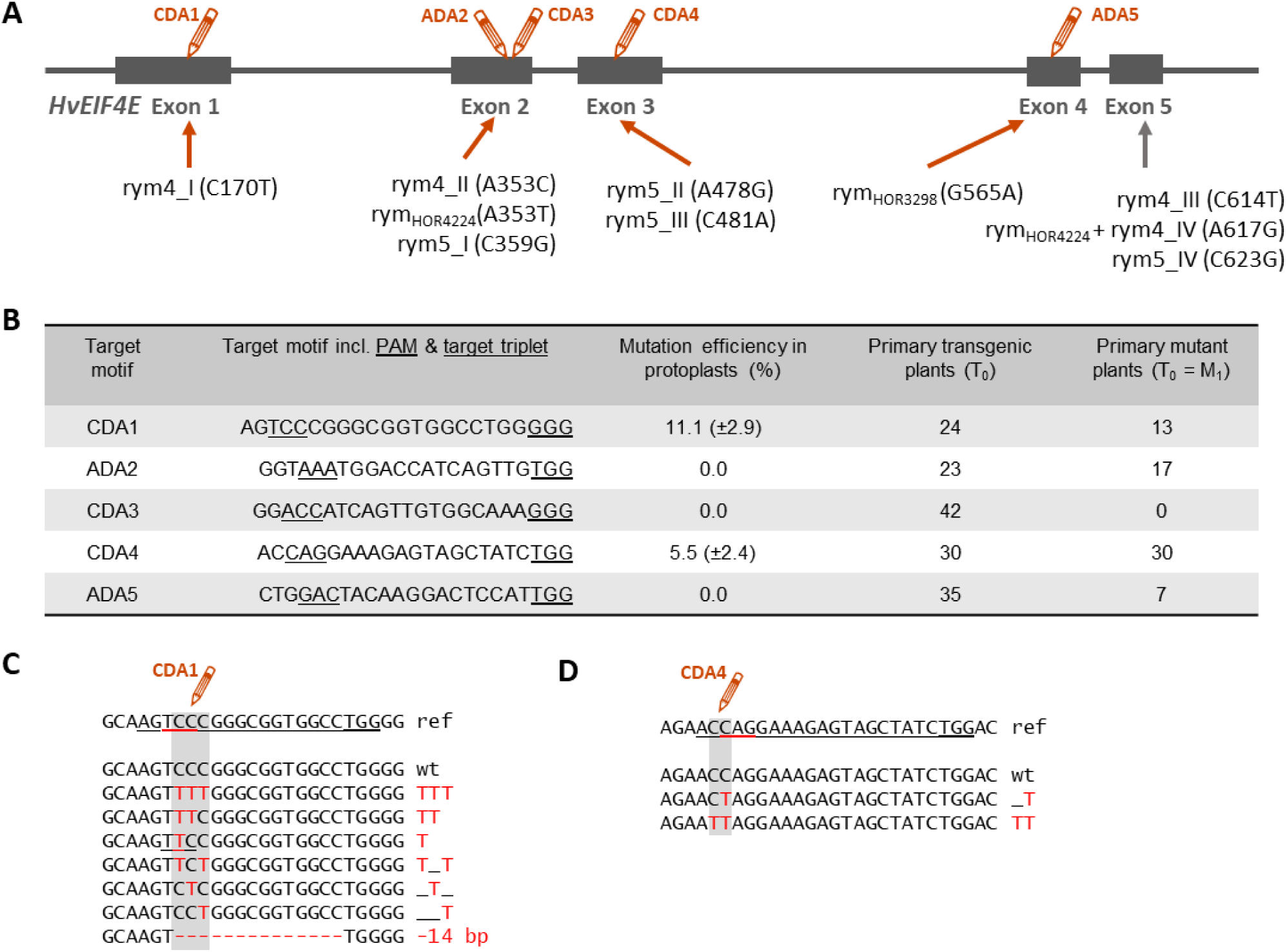
Target motifs and mutation efficiency; Mutation patterns in barley mesophyll protoplasts, A: Gene structure of the *Hordeum vulgare EIF4E* gene with locations of the chosen target motifs for base editing in regions with nucleotide polymorphisms previously described for the resistance-conferring alleles *rym4*, *rym5*, *rym_HOR3298_ and rym_HOR4224_*. Pencils represent base editing targets, CDA: Cytidine deaminase-nCas9 fusion, ADA: Adenine deaminase-nCas9 fusion, grey line represents gene introns, grey boxes represent exons. **B:** Summary of target motif sequences including the respective protospacer adjacent motif (PAM, double underlined) and the coding triplet addressed by base editing (single-underlined). Mutation efficiency in protoplast assay and primary mutants based on amplicon deep-sequencing. Primary transgenic plants from *Agrobacterium*-mediated DNA transfer to microspore-derived microcalli of winter barley cv. ’Igri’. **C + D:** Results of amplicon deep-sequencing after PEG-mediated transformation of protoplasts with intermediate vectors carrying CDA-nCas9 fusion and respective gRNA for target motifs CDA1 (**C**) and CDA4 (**D**). Cytidine bases in editing windows are marked by grey box, base edits indicated by red letters, deletions by red dashes, ref: reference sequence with the target motif underlined, the PAM double-underlined and the targeted coding triplet double-underlined in red, wt: wild- type sequence.

### Vector cloning

Vectors were cloned using the CasCADE vector system (Hoffie et al., in preparation). A generic module for gRNA position 1 with the *U3* promotor from *Oryza sativa* was used (pIK1). Target-specific sequences of the gRNA were ordered as DNA oligonucleotides (see Supplementary Table S1) with specific overhangs for BsaI-based cloning into pIK1. Golden Gate assembly of gRNA and Cas modules with nCas9-CDA (pSH272, D10A-nCas9 with C-terminal fusion of PmCDA1 according to Shimatani et al. (2017)) and nCas9-ADA (pSH273, D10A-nCas9 with N-terminal fusion of EcTadA-EcTadA* according to Li et al. (2018)) fusions, driven by the *Zea mays Polyubiquitin 1* (*ZmUbi1*) promotor, was performed according to the CasCADE protocol to generate the following vectors: pRH20 (EIF4E_CDA1), pRH21 (EIF4E_ADA2), pRH22 (EIF4E_CDA3), pRH23 (EIF4E_CDA4) and pME25 (EIF4E_ADA5). The gRNA and nCas9-CDA/ -ADA expression units were cloned via SfiI into the binary vector p6i-2x35S-TE9 (DNA Cloning Service, Hamburg, Germany) that harbours an *hpt* gene under control of a doubled-enhanced *CaMV35S* promoter in its transfer-DNA for plant selection, resulting in the respective binary vectors for *Agrobacterium*-mediated barley transformation pRH27, pRH28, pRH29, pRH30 and pME35.

### Pre-validation of constructs via transient expression in protoplasts

The intermediate vectors were first tested by transient expression in mesophyll protoplasts isolated from leaves of one week-old, etiolated seedlings of winter barley cv. ’Igri’ based on the protocol of Shan et al. (2014). In brief, thirty leaves of barley seedlings were chopped into small fragments using a razor blade, and cell walls were digested with Macerozyme R-10 and cellulase R-10 (DUCHEFA BIOCHEMIE B.V, Haarlem/Netherlands). Protoplasts were purified by sieving, and after PEG-mediated transformation with pRH20, pRH21, pRH22, pRH23 and pME25, they were incubated for 60 h at 21 °C in the dark. All protoplast transformations were conducted in three replications. To check for transformation efficiency, a *GFP* expression construct was used to transform a control sample of protoplasts. After incubation, the GFP-expressing portion of protoplasts of this control sample was determined using an epifluorescence microscope (AX200M, Zeiss, Oberkochen/Germany), and DNA was extracted from the other protoplast samples transformed using the intermediate *cas9*/gRNA constructs. Around 150 bp of the target regions were amplified using specific primers (see Supplementary Table S1), followed by deep-sequencing of amplicons, which was performed by a commercial service provider on an Illumina MiSeq platform (Amplicon-EZ service by Genewiz, Leipzig/ Germany). Mutation efficiencies were calculated individually for each replicate as proportion of sequencing reads with mutation in relation to the total number of reads including those with the wild-type sequence.

### Agrobacterium-mediated transformation using barley microspore-derived plant regeneration

The binary vectors pRH27, pRH28, pRH29, pRH30 and pME35 were transfected by electroporation into the *Agrobacterium* strain LBA4404 harbouring the hypervirulence- conferring plasmid pSB1 for DNA transfer to barley microspore-derived microcalli (Kumlehn et al., 2006). In brief, pre-mitotic (highly vacuolated) microspores were isolated from winter barley cv. ’Igri’. After microspore cultivation for 1 week to initiate cell proliferation, resultant multicellular pollen was subjected to DNA transfer by co-cultivation with *Agrobacterium* carrying the binary vectors as mentioned above. Upon further cultivation under selective conditions, transgenic plantlets were generated from calli or embryogenic structures as previously described in detail by Kumlehn et al. (2006) and Hoffie et al. (2021a).

### Genotyping of primary transgenic and mutant plants

Leaf samples were taken from putative primary transgenic plants and DNA was extracted and analysed for the presence of the *cas9* transgene by PCR using specific primers (see Supplementary Table S1). The plants were screened for mutations by PCR amplification of the target regions using specific primers (see Supplementary Table S1) followed by amplicon deep-sequencing by a commercial provider using the Illumina MiSeq platform. The mutagenesis efficiency was calculated based on the mutated sequencing reads in relation to the total number of sequencing reads. The ploidy level of all regenerants was measured by flow cytometry to identify spontaneous doubled-haploid plants. Plants that were found to be still haploid were subjected to colchicine treatment for genome duplication, according to Hoffie et al. (2021a). Mutant plants were grown to maturity after self-pollination and grains were harvested. To analyse the M_2_ generation, progeny comprising 12 or 24 individuals were sown from selected primary mutants. DNA was extracted from leaf samples at early seedling stage and the inheritance of the transgene was screened by PCR using *cas9-* and *hpt*-specific primers (see Supplementary Table S1), and the target regions were PCR-amplified and screened by Sanger sequencing for induced base substitutions.

### Phenotyping: BaMMV inoculation and resistance screening by DAS-ELISA, yield data

To screen base-edited lines for Bymovirus resistance, plants were mechanically inoculated with BaMMV according to a protocol of Habekuss et al. (2008) and Wang et al. (2021a). For the first screening, 15 plants per M_3_ line were grown each, resulting in a total of 160 edited seedlings cultivated along with 24 Igri wild-type plants in a growth chamber at 12 °C and 16 h photoperiod and inoculated twice with leaf sap of BaMMV-ASL-infected barley plants with an interval of five to seven days at the three-leaf stage. Six to eight weeks after the first inoculation, visible symptoms were assessed. Additionally, the plants were screened for virus particles by double-antibody sandwich enzyme-linked immunosorbent assay (DAS-ELISA). Plants of the susceptible genotypes ‘Maris Otter’ and ‘Uschi’ served as controls for infection and phenotyping. Resistance or susceptibility was evaluated based on the extinction *E*_405_ in relation to positive and negative controls. Simultaneously, DNA was extracted from leaf samples of all test plants, PCR was performed for the *cas9* transgene and the target region was amplified and Sanger sequenced as described above. In a second batch, in total 63 plants of selected M_4_ lines were phenotyped and genotyped as described above.

After resistance screening, M_3_ and M_4_ plants were cultivated in a glasshouse with 18-20 °C/ 12- 14 °C day/night with 16 h light to maturity, harvested and ears were threshed. The total grain number of each individual plant was counted and total grain weight was measured. Thousand-grain weight (TGW) was calculated as:

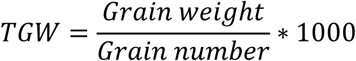

### Statistical analysis

Qualitative resistance data was analysed by a global Fisher’s Exact test due to variation in sample size between genotypes, followed by post-hoc pairwise Fisher’s Exact test with p adjustment according to Benjamin-Hochberg.

Quantitative yield data was analysed by Shapiro-Wilk test for normal distribution and by Brown-Forsythe test for equal variance to decide about subsequent data processing. Based on that, significant differences in grain number were identified by Welch-ANOVA and Games-Howell post-hoc test (normal distribution but high differences in variance). Data of thousand-grain weight was further analysed by Kruskal-Wallis rank sum test, followed by Dunn’s Post-Hoc test with p adjustment according to Benjamin-Hochberg (no normal distribution, high differences in variance).

All tests were performed using the statistics software R version 4.6.0 (R Core Team, 2026). Plots were generated with R package ggplot2 (Wickham, 2016). R code was optimised with support of Google’s Gemma 4 31B Instruct large language model, locally hosted and provided by ChatAI (GWDG, Göttingen, Germany, Doosthosseini et al. (2026)). Full R code and results can be found in the supplements S2.

## Results

### Pre-validation of transformation constructs in protoplasts

After the intermediate (non-binary) vectors pRH20, pRH21, pRH22, pRH23 and pME25 had been generated using the CasCADE system, they were tested in protoplasts to pre-validate their functionality and mutagenesis efficiency by deep sequencing of PCR amplicons of the target regions. A general overview is given in Figure 1B. Mutagenesis efficiencies for CDA targets reached up to 11 %, while no mutations were detected in both ADA-addressed targets. For the CDA1 and CDA4 targets, Figures 1C and 1D show the sequence variants obtained from protoplasts. The editing window of both target motifs contained several Cs that were converted individually or in different combinations into Ts. This resulted in a number of new alleles that would also lead to a variety of amino acid changes after translation.

### Screening of primary mutants and their progeny for base edits

All five binary vectors pRH27, pRH28, pRH29, pRH30 and pME35 were used for stable transformation by *Agrobacterium*-mediated DNA transfer in microspore cultures of the winter barley cultivar ’Igri’.

Several transformation experiments were carried out in each case, regenerating a total of 23 to 42 plants for each construct (see Figure 1B). For all target motifs except CDA3, plants were identified by deep sequencing of PCR amplicons that had a higher proportion of sequencing reads with alterations than the wild-type control (see summary in Figure 1B). The regenerated plants were mostly chimeric, which could be inferred from the large number of different alleles, as exemplified for three target motifs in Figure 2. In the target motif CDA1, there are three Cs in a row within the editing window. In plant BPR42/1P3, a total of 61 % of the sequencing reads had a base substitution in the target region (for comparison: ’Igri’ wild-type had 1 % only). Of these, one mutation accounted for almost 50 %, suggesting that it was a heterozygous allele. Here, a C was replaced by a T, which changed the affected coding triplet from CGG (arginine) to TGG (tryptophan). In the same plant, other alleles occurred with much lower frequency, suggesting that these were confined to smaller sectors of the plant. In total, three different base edits were identified in this plant in addition to the wild-type allele, two of which led to changes in the resulting amino acid. The change from serine (TCC) to phenylalanine (TTC), as described for the allele *rym4* at this position, is also among the edited sequence variants.

**Figure 2:**
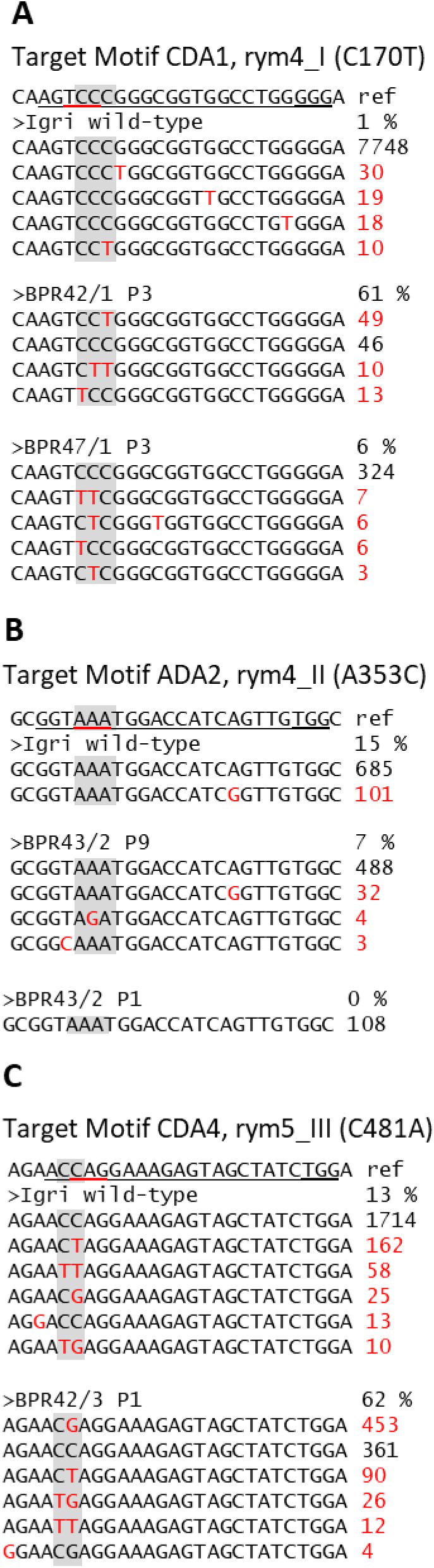
Mutation patterns in primary mutant barley plants, Results of amplicon deep- sequencing after *Agrobacterium*-mediated DNA transfer to microspore-derived microcalli of winter barley cv. ’Igri’ with binary vectors carrying a CDA-nCas9 or ADA-nCas9 fusion and respective gRNA for target motifs CDA1 (**A**), ADA2 (**B**) and CDA4 (**C**). Cytidine or adenine bases in editing windows are marked by grey box, base edits indicated by red letters, deletions by red dashes, ref: reference sequence with the target motif underlined and the PAM double-underlined, wt: wild-type sequence, numbers indicate the number of reads of the respective sequence contig.

In the target region ADA2, the proportion of A-to-G-substitutions in the target region was rather low (max. 7 %), but in contrast to the protoplast assay, at least a few of such modifications were found at the plant level. In plant BPR43/2P9, one allele changed in the target triplet was found with very low frequency, which was edited from AAA (lysine) to AGA (glycine). The substitution described for *rym4* at this position is ACA, which entails a change of the encoded lysine (wild-type) to threonine (*rym4_II*). At the CDA4 target motif, C-to-T-changes occurred with a relatively high frequency of 13 % in the wild-type sample, most likely due to cross-reactions within the pooled samples during library preparation. However, the mutation frequency in base editing plants reached 62 % in case of BPR43/2P1 with a C-to-G-substitution in the editing window being the most frequent edit.

Among the primary mutants, those with comparatively high proportions of mutant sequencing reads were selected after maturity and M_2_ progenies of 12 or 24 plants were grown, i.e. four progenies per target region CDA1, ADA2 and ADA4, and five progenies for target region CDA4 with a total of 84 plants for target region CDA1 and a total of 96 plants each per target region ADA2, CDA4 and ADA4, respectively. After DNA extraction, the target regions were PCR-amplified and analysed for the inheritance of the induced base edits by Sanger sequencing. In parallel, T-DNA-specific PCR was used to follow the segregation of the transgene. The results of all plants are shown in Table S2. No mutations were detected in the progeny at any of the ADA target motifs. In contrast, both heterozygous and homozygous mutants were identified for the CDA1 and CDA4 target motifs. The proportion of mutants among siblings ranged from 0 to 100 %. The CDA1 target motif has three Cs within the editing window, distributed between the two triplets TCC (serine, pos. 57) and CGG (arginine, pos. 58). In the four M_2_ families studied, the mutation TCC to TTC (C170T) occurred most frequently (see Figure 3A), resulting in an amino acid substitution from serine to phenylalanine (S57F), i.e. exactly the change that is also characteristic for *rym4* at this position. Base edits at the second triplet from CGG to TGG (C172T) replace arginine with trypsin (R58W). In two M_2_ plants, C-to-T edits outside the expected editing window were also found, resulting in amino acid changes at protein positions 59 and 61 (A59V, A61V). Several plants with a C-to-G-edit (C170G; S57C) also occurred in the progenies of BPR42/1_P3 and BPR44/1_P1. In total, different combinations of the edited bases and resulting amino acid substitutions generated seven different alleles at the CDA1 target region, shown in Figure 3A and 3C. In the majority of mutated M_2_ plants, the respective mutation was already homozygous.

**Figure 3:**
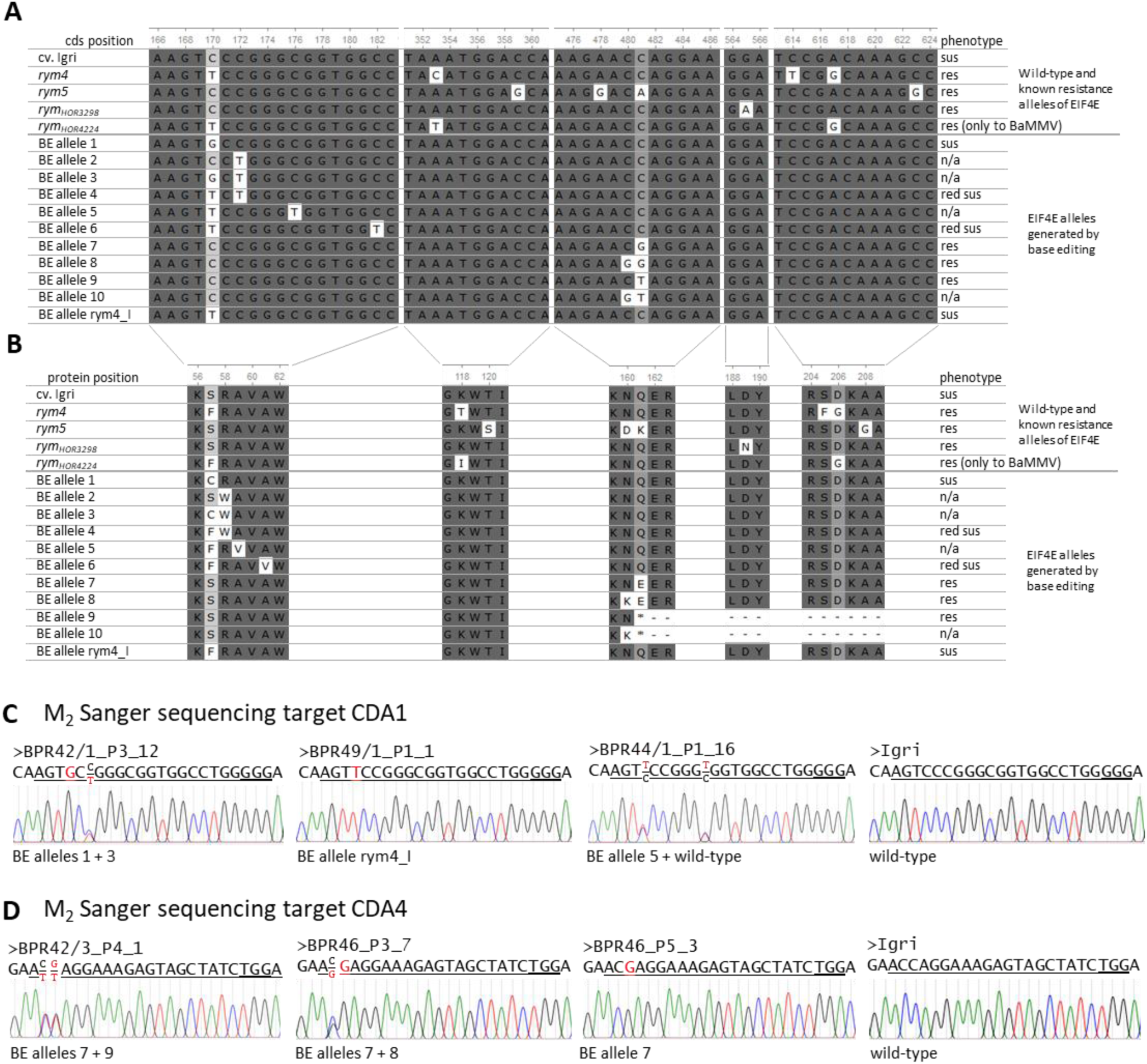
***EIF4E* gene sequence variation and resulting protein sequence variation inherited to M_2_ plants,** Sequences generated by base editing at targets CDA1 and CDA4, inherited to M2 generation in comparison to genetic variation previously discovered: **A:** Alignment of five sections of the *EIF4E* coding sequence (cds) of all base-edited (BE) alleles to the wild-type sequence of the susceptible winter barley cv. Igri as well as the resistance alleles *rym4*, *rym5*, *rym_HOR3298_* and *rym_HOR4224_*. **B:** Protein sequence variation (amino acid one letter code) resulting from genetic variation of the same sections of the cds. Gray scale visualises sequence similarity; cds and protein position of nucleobases and amino acid residues are given above the respective alignment; Phenotype: sus: susceptible, res: resistant, red sus: reduced susceptibility to BaMMV/ BaYMV infection; *: stop codon. **C + D**: Sanger chromatograms from selected, representative M_2_ barley plants compared to wild-type for target motif CDA1 (**C**) and target motif CDA4 (**D**). Double peaks indicate heterozygous base edits (both bases shown as fraction). Sequence of target motif underlined, PAM double-underlined, red letters indicate mutations.

The CDA4 target motif has two Cs within the editing window that belong to two triplets: AAC (asparagine, pos. 160) and CAG (glutamine, pos. 161). Plants with base edits were identified in four of the five M_2_ families examined. Different combinations of C-to-T- and C-to-G-substitutions occurred. Since both triplets were affected, there was also a corresponding variability in amino acid changes. For the first codon, the change AAC to AAT (C480T) occurred most frequently, whereby it is a "wobble" base, so that this triplet likewise codes for asparagine. The substitution AAC to AAG (C480G) occurred less frequently but codes for lysine (N160K). At the second triplet, the substitution CAG to GAG (C481G), which codes for glutamate (Glu), and CAG to TAG (C481T), which leads to the formation of a stop codon (Q161*), occurred with similar frequency. These modifications of the two Cs occurred individually or in combination in the analysed offspring, resulting in a total of four novel alleles. However, the substitutions of asparagine by aspartic acid (N160D) and glutamine by lysine (Q161K), which are characteristic for the *rym5* allele at this position, were not among the variants generated.

In total, this base editing approach in the susceptible winter barley ’Igri’ resulted in a polymorphism described for the resistance-mediating allele *rym4* as well as in 10 further, novel alleles that lead to amino acid changes at relevant positions in the eIF4E protein (see Figure 3A and full coding and protein sequences in S1).

### BaMMV inoculation of base-edited plants

Based on the previous genotypic screening, in total 12 M_3_ lines were selected for infection tests with BaMMV aiming to cover all base-edited (BE) alleles. For each line, 15 grains were sown and 13 to 15 plants were obtained, resulting in a total of 160 base-edited plants that were inoculated with BaMMV. Due to segregation within the tested M_3_ families, the number of plants was unequally distributed between all eleven BE alleles, but a minimum of eight plants was tested for each generated *EIF4E* sequence variant. After infection, virus-induced symptoms were assessed (Figure 4B) and presence of virus particles was tested by ELISA. As control, 24 Igri wild-type plants were inoculated in parallel, out of which 18 (75 %) were successfully infected with BaMMV. At the CDA1 target, in total seven different alleles were generated, including a sequence variant that occurred, beside three others, as part of the biologically evolved resistance conferring *rym4* allele of *EIF4E*. Remarkably, out of 38 homozygous plants with this rym4_I allele (C170T; S57F), 35 (92 %) were susceptible to the BaMMV infection. Homozygous plants of BE allele 1 (C170G; S57C) and BE allele 6 (C170T + C182T; S57F + A61V) were also susceptible to BaMMV infection. Five out of eight plants (63 %) carrying the BE allele 4 (C170T + C172T; S57F + R58W) were proven virus-free by ELISA. For BE alleles 2 (C172T, R58W) and 5 (C170T + C176T, S57F + A59V) no homozygous plants were obtained, whereas for BE allele 3 (C170G + C172T, S57C + R58W) only two.

**Figure 4:**
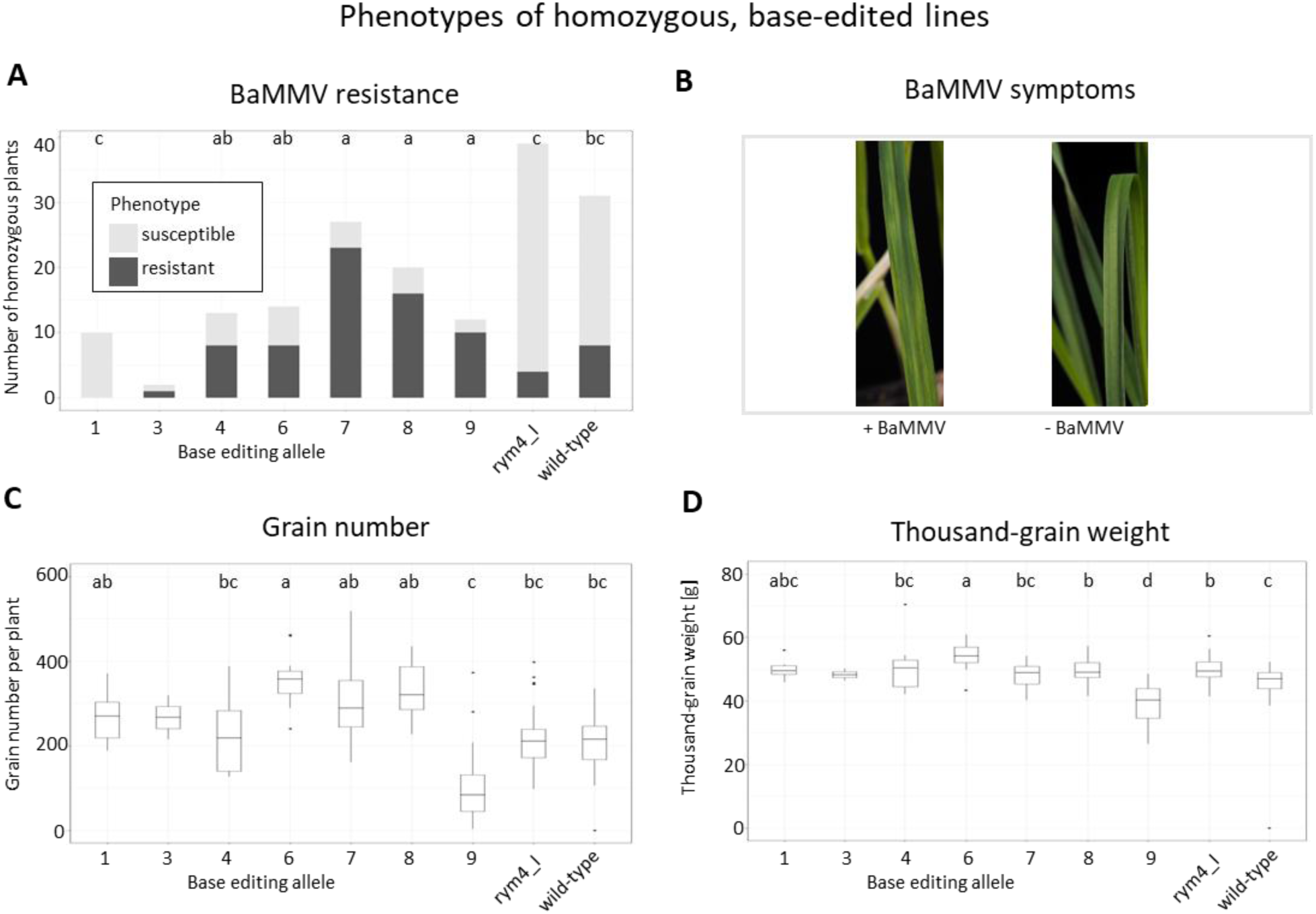
Phenotypes of homozygous, base edited M_3_ and M_4_ lines, A: total number of homozygous, resistant (dark gray) or susceptible (light gray) plants per *EIF4E* allele after mechanical inoculation with BaMMV, based on ELISA data. Exclusively different letters indicate significant differences (p≤0.001) based on pairwise Fisher’s Exact test with p-adjustment according to Benjamin-Hochberg. **B:** Visual symptoms of BaMMV infected barley leaves in comparison to an uninfected plant. **C + D:** Grain number per plant (**C**) and thousand-grain weight (**D**) were assessed as yield parameters per individual plant and shown as box plots allele-wise. Exclusively different letters indicate significant differences (p≤0.03). Statistical analysis of grain number was performed by Shapiro-Wilk test for normal distribution, Brown-Forsythe test for equal variance, followed by Welch-ANOVA and pairwise Games-Howell post-hoc test. Thousand- grain weight data was analysed by Shapiro-Wilk test for normal distribution, Brown-Forsythe test for equal variance, followed by Kruskal-Wallis test and Dunn’s post-hoc test with p-adjustment according to Benjamin-Hochberg. All data in this figure are based on in total 168 homozygous plants, BE allele 3 was excluded from pairwise comparisons due to low sample size (n=2).

Base editing of target CDA4 generated four new alleles of the gene. Seven out of eleven plants (64 %) homozygous for BE allele 7 (C481G; Q161E) were not infected after BaMMV inoculation. Notably, all seven plants that were biallelic for BE alleles 7 and 8 (C480G + C481G; N160K + Q161E) and all seven plants that were biallelic for BE alleles 7 and 9 (C481T; Q161*) were resistant to infection with the virus. BE allele 10 (C480G + C481T; N160K + Q161*) was not inherited to the M_3_ generation and no plants were obtained for infection tests. All plant data of M_3_ are listed in Table S3.

To further investigate the effect of *EIF4E* base edits on BaMMV susceptibility, a second batch of plants was subject to BaMMV infection tests. Selected lines with BE alleles rym4_I, 4, 6, 7, 8 and 9, in total 63 plants, were grown and infected. Especially for the alleles generated at the CDA4 target showed an effect: 15 out of 16 plants homozygous for the BE allele 7 and 15 out of 17 plants homozygous for BE allele 8 were resistant to the BaMMV infection (complete data see Table S3). To summarise the effect of base edits within the *EIF4E* gene, all data from infection tests were counted together for homozygously edited plants (Figure 4A). A global Fisher’s Exact test revealed significant differences between the different genotypes (p = 0.0005). Pairwise Fisher’s Exact tests between all genotypes identified three groups: BE alleles 7, 8 and 9 (a) exhibit high resistance rates that are significantly (p < 0.001) different from BE allele 1, rym4_I and wild-type (c) with very high susceptibility. BE alleles 4 and 6 confer significantly stronger resistance than BE allele 1 and rym4_I (p ≤ 0.001) but do not differ significantly from BE alleles 7, 8 and 9 (p > 0.1). BE allele 3 was excluded from statistical analysis due to the too low number of plants (2), while for the other genotypes a minimum of ten plants was analysed.

### Yield-related traits affected by base editing

All plants were grown to maturity and the yield-relevant traits grain number per plant and total grain weight were assessed and the thousand-grain weight was calculated. Again, the data was summarised for all the homozygous plants per genotype (Figure 4C and 4D). For grain number, a Welch-ANOVA (normal distribution but unequal variance between genotypes) revealed significant differences between genotypes (p = 6.3*10^-6^). Pair-wise comparison by Games-Howell post-hoc test identified individual differences between genotypes: Plants with BE allele 9 (knockout owing to premature stop-codon) exhibit a significantly lower grain number than plants with BE alleles 1, 6, 7 and 8 (p ≤ 0.02), while it was weaker, yet not significantly different from plants with rym4_I and wild-type alleles (p > 0.1).

Thousand grain weight data (Figure 4D) was analysed by Kruskal-Wallis rank sum test revealed significant differences between genotypes (p = 5.9*10^-8^). Pairwise comparison by Dunn’s post-hoc test proved BE allele 9 to result in a significantly lower TGW than BE alleles 1, 4, 6, 7, 8 and rym4_I as well as the wild-type (p ≤ 0.03), while BE alleles 1, 6, 8 and rym4_I are causing significantly higher TGW than wild-type plants (p ≤ 0.03). TGW of plants with BE allele 7 was on a par with wild-type plants (p = 0.29).

Taken together, base editing alleles 7 and 8 combine high resistance against BaMMV infection by maintaining high yield potential.

## Discussion

### Base editing in barley protoplasts and plants

Base editing constructs were created for five *rym4*, *rym5, rym_HOR4224_* or *rym_HOR3298_* polymorphic positions in the *HvEIF4E* gene. Base editing was successfully performed in two of these target motifs, thus generating a number of novel alleles. Two of the target motifs, where detectable mutations could be induced neither in protoplasts nor in stably transformed plants, i.e. ADA2 and CDA3, are located close to each other in the second exon. In previous work, mutations were not obtained here either, even using a classical approach of targeted mutagenesis, so that it is possibly a position effect in the genomic context that makes this target region difficult to access for Cas9 (Hoffie et al., 2021b). In addition, the approach of mutating very specific positions in the target gene is associated with a lack of flexibility to select targets with particularly good *in silico* predicted scores for high mutagenesis efficiency.

While relatively high mutation rates were achieved at the target motifs CDA1 and CDA4 and were also inherited to the next generations (as followed until M_4_), no base edits were found in the M_2_ at the second adenine deaminase target motif ADA5. The sequence variants found in primary transgenics were most likely amplification and/or sequencing errors, as they also occurred at similarly low frequencies in the wild-type controls. Thus, based on the very limited number of two targets tested, there is no evidence of the functionality adenosine deaminase base editors in barley. The module used in the CasCADE system adapted the ABE7.10 fusion protein of the ecTadA wild-type and the derivative ecTadA* developed by directed evolution by Gaudelli et al. (2017). In mammalian cells, editing efficiencies of up to 70 % were achieved with this ABE variant. In *Arabidopsis* and oilseed rape protoplasts, Kang et al. (2018) were still able to achieve up to 4 % and up to 9 % A-to-G substitutions, respectively, in this configuration; in stably transformed plants, up to 85 % of M_2_ progeny showed mutation rates of ≥50 %. At the ADA target motifs in this study, no A-to-G substitutions occurred in protoplasts and only in a few stable plants in a very low proportion. In M_2_, none with corresponding base edits were found among a total of 192 plants. Gaillochet et al. (2023) optimised nCas9 and nCas12a base editors for wheat and barley and observed editing efficiencies with an enhanced nCas9-TadA-fusion (version 8e) of up to 22 % on an extrachromosomal GFP target in protoplasts. However, even this optimised base editor was only effective at one out of four endogenous targets in wheat, underlining the high variation of editing efficiency between individual targets.

Recently, Zhang et al. (2025) developed an entirely new cytosine base editor based on the fusion of an engineered bacterial SsdA (single-strand DNA deaminase toxin A) to nCas9 and proved it efficient base editing activity in barley and rice protoplasts and rice plants.

The protoplast assay proved to be a reliable approach for base editing to predict the functionality and efficiency of gene editing constructs although the full cycle of DNA repair required for base editing includes DNA replication (Nambiar et al., 2022). The results from protoplasts correlated clearly with those in stably transformed plants. In each case, base editing was most efficient at the target motif CDA1, while the efficiency at CDA4 was slightly lower, and no changes could be detected at ADA2, CDA3 and ADA5 that could be convincingly distinguished from sequencing errors.

At cytidine deaminase targets, numerous C-to-G changes were found in plants in addition to the expected C-to-T substitutions. At target motif CDA4, these accounted for about one third of the observed base edits in M_2_ and led to additional new alleles. A similar proportion was also described by Nishida et al. (2016) in their first experiments with *Pm*CDA1 base editors. In our case, additional variation was useful and also for other approaches this will broaden the scope for certain base substitutions. We targeted Cs in a range from position 2 to 5 of the target motif (considering the most distal base from the PAM as base position 1). However, we observed rare base editing events until position 16 of the target motif (BE allele 6), making the editing window wider than expected. The editing window is mainly influenced by length and rigidity of the linker between nCas9 and cytidine deaminase. We used the XTEN linker as previously described by Komor et al. (2016) with a presumed editing window of five bases, peaking around position 7 of the target motif. Distal edits as at position 16 in our case were not reported. Shi et al. (2026) recently investigated the effect of different linker length and properties for base editing. The editing window was varying between six and 14 bases, always peaking around bases 5 to 7 auf the protospacer, even with very long and flexible linkers. This underlines that our BE allele 6 with an edit at position 16 is very rare and indeed it goes back to only one single event found in the M_2_ mutant plant BPR49/1_P3-16.

Inheritance of induced base edits was followed until the M_4_ generation and was proven also for transgene-free lines. For instance, family BPR42/3P4-7-12 was PCR negative for the transgene and exhibited a Mendelian inheritance pattern of 3:5:2 for homozygous BE allele 7: heterozygous 7/9: homozygous BE allele 9. Since both alleles confer resistance to BaMMV, no segregation of the phenotype was observed though. Due to transformation of haploid microspores followed by spontaneous or induced genome duplication, most lines were homozygous for the T-DNA and no segregation of transgene and induced mutations was observed. In such cases, backcrosses with Igri wild-type are required to dissect transgene and base edits. The occurrence of transgene-free segregants, despite the methodological context of generating doubled haploids, can be explained by the fact that editing events as well as T-DNA integration can also take place after spontaneous genome duplication, resulting in heterozygous mutations and hemizygous DNA integration in diploid cells.

### Virus inoculation reveals resistance conferring base-edited alleles

In M_3_ and M_4_, seven base editing alleles were tested along the re-created rym4_I allele and wild-type plants for BaMMV resistance. While plants carrying BE alleles 1 and 3 were as highly susceptible as plants with the rym4_I allele and wild-type plants, the susceptibility was reduced in plants with BE alleles 4 and 6, while plants with BE alleles 7, 8 and 9 can be considered resistant. For BE allele 9, a pre-mature stop codon leading to a loss-of-function allele, this is in line with the knock-out of *HvEIF4E* that was previously described in barley, wheat and other plant species (Hoffie et al., 2021b; Kan et al., 2023; Schmitt-Keichinger, 2019). As the frame-shift knockout lines, also BE allele 9 plants were strongly impaired in grain setting and filling, underlining the importance of a functional *EIF4E* gene for proper plant performance.

BE alleles 7 and 8 (both generated at target CDA4) combined BaMMV resistance as high as in BE allele 9 plants while maintaining yield potential as high as in wild-type plants, making these two alleles the most interesting ones for breeding approaches. The direct comparison of the phenotypes of the stop codon BE allele 9 and the frame-shift knockout lines from our previous work strongly indicate that BE alleles 7 and 8 maintain the functionality of eIF4E in protein biosynthesis, although direct binding of edited eIF4E proteins to the m7G-cap of mRNA was not studied here.

The BaMMV-ASL isolate was chosen thanks to its proven suitability for reproducible mechanical inoculation. However, it is not virulent on *rym4* and *rym5*. Trials on infested fields with natural root infection by the vector *Polymyxa graminis* and with different BaMMV and BaYMV isolates are necessary to further validate the suitability of BE alleles for breeding.

By comparing several alleles of *EIF4E* in barley (see Figure 3A), our base editing alleles indeed contribute new variation to this gene, that exhibits already particularly high diversity due to strong selection pressure by co-evolution with bymoviruses (Hofinger et al., 2011). Our results also coincide with the screen of more than 40 *EIF4E* haplotypes in barley by Hofinger et al. (2011) since they also found particular high variation at coding sequence positions 480 to 483 of the coding sequence. This underlines the importance of the coded amino acid residues around protein position 160 for interaction with potyviruses in evolution.

The resistance mechanism is well described for eIF4E of several plants and their potyviruses: Sequence alteration at the binding domains of the eIF4E protein inhibit the interaction with the viral cap-like VpG-protein and avoid translation of the viral RNA (Coutinho de Oliveira et al., 2019). Since we targeted previously described positions within the *EIF4E* gene in barley, we can assume that the base edits in BE alleles 7 and 8 have the same effect. Both these alleles are altered at the CDA4 target, creating changes of amino acid residues 160 and/or 161 (N160K/ Q161E). That is in line with the finding that especially the residues 158 and 163 of the barley eIF4E are involved in binding of the VpG cap (Li et al., 2016). Remarkably, the re-creation of the rym4_I polymorphism at the target CDA1 (S57F) had no effect on susceptibility. While the *rym4* allele also carries additional mutations (see Figure 3A), the one that was re-created here does not contribute to the resistance or only in combination with the other polymorphisms. This needs to be further investigated, while our results suggest that not all mutations of the *rym4* and *rym5* alleles contribute equally to the resistance against bymoviruses.

### Genome editing of EIF4E beyond knockouts

Wen et al. (2024) generated a combination of knockout and in-frame deletion mutants of *EIF4E1* and *EIF4E2* in maize that were linked to potyvirus resistance using a classical Cas9-mediated genome editing approach. Bastet et al. (2019) dissected multiple polymorphisms found in *Pisum sativum EIF4E* and their individual effect on potyvirus resistance in *Arabidopsis*. Also, in that pathosystem, not all spontaneous variations found in *PsEIF4E* contributed to the resistant phenotype. Furthermore, these authors successfully used base editing to re-create one of these polymorphisms associated with resistance by introducing a C-to-G change in the *Arabidopsis EIF4E* gene. The same lab later applied base editing to tomato *EIF4E1* and generated two base editing alleles that conferred (partial) resistance to different potyviruses (Kuroiwa et al., 2023). Our study represents the first report of new allelic variation generated in *EIF4E* by base editing in a monocotyledonous crop plant and the first base editing approach in barley plants at all.

With regard to the *rym4* and *rym5* alleles, that have been widely used in winter barley breeding in Europe, it is well described that selection pressure leads to evolution of new bymovirus strains that have overcome these resistance alleles already. Stacking of different alleles and combining different resistance mechanisms can be a promising approach to prevent that. In regards to bymoviruses in barley, it was previously shown that the full knockout of the susceptibility factor *HvPDIL5-1* leads to full resistance of barley without adverse effects on yield traits (Cheng et al., 2022; Hoffie et al., 2023). Consequently, combing the knockout of *PDIL5-1* with base editing of *EIF4E* is a promising approach to generate a long-lasting potyvirus resistance in barley and other crop plants.

### Summary and outlook

In summary, our study demonstrates the potential of base editing, especially when employing an nCas9-cytidine deaminase fusion to generate novel allelic variants in specifically addressed target motifs as well to exactly reproduce valuable alleles known from other genotypic contexts. However, the latter – that is, the precise substitution of one specific nucleobase for another – remains a considerable challenge even with this approach. In particular, the ability to precisely and effectively target the position to be changed is limited by the need for a suitably located PAM and efficiency-determining features of the gRNA, as well as by the fact that CDA and ADA have so far been largely restricted to C-to-T and A-to-G changes. Improved possibilities in the precise addressing of specific genome positions are offered by fusions from CDA or ADA with Cas derivatives that have greater flexibility in terms of PAMs, such as nCas-NG, which can use all NG motifs as PAMs instead of NGG. On the other hand, these PAM-relaxed endonucleases are also associated with reduced editing efficiency (Nishimasu et al., 2018). Base editing primarily offers the possibility of generating new allelic diversity beyond knockouts at predetermined positions in the target genes. In the present study, the *HvEIF4E* gene was chosen as a target of great relevance from a breeding perspective, with a total of 10 novel alleles being generated. The BE alleles 1, 3 and the re-created rym4_I allele had no detectable effect on resistance, while the loss-of-function BE allele 9 (stop codon) lead to plant resistance against BaMMV infection, which, however, was at the cost of a strongly reduced number of grains per plant and lower thousand-grain weight. In contrast, BE alleles 7 and 8 combined resistance to BaMMV infections with yield parameters on a par with wild-type plants, proving the potential of base editing of *EIF4E* for resistance breeding. Given the good knowledge about *EIF4E* and potyviruses in many plant species, this will also be a promising approach for resistance breeding in other mono- and dicotyledonous plants.

## Supporting information

S1 sequences

S2 R code and output

Table S1 oligos

Table S2 M2 plant data genotype

Table S3 plant data M3 + M4 genotype + phenotype

## Author Contributions

JK and RH conceived the project and acquired the funding. RH devised the experimental setup. RH and IH performed the experiments, AH and DP supervised the BaMMV infection tests, RH analysed the data. RH wrote the manuscript. All authors edited and approved the manuscript.

## Funding

The research was funded in frame of the IdeMoDeResBar project 031B0887 (phase 2) by the German Federal Ministry of Education and Science (BMBF) as well as by basic funding of IPK and JKI.

## Conflict of Interest

The authors declare that the research was conducted in the absence of any commercial or financial relationships that could be construed as a potential conflict of interest.

## Acknowledgments

The authors thank Ingrid Otto and Carola Bollmann (barley transformation and protoplast assay), Katy Niedung and Ingrid Dubsky (BaMMV infection tests), Claudia Baumann and Markus Erhardt (DNA extraction, PCR and sample preparation for sequencing) for excellent technical assistance.

## Supplementary Material

Sequence S1: *EIF4E* allelic coding sequences and resulting protein sequences

R code S2: full R code and results used for statistical analysis

Table S1: Oligonucleotide sequences

Table S2: M_2_ plant data: genotype

Table S3: M_3_ and M_4_ plant data: genotype and phenotype

## Notes

### Competing Interest Statement

The authors have declared no competing interest.

