## Supplementary material for "Base editing of *EIF4E* creates novel resistance alleles against bymoviruses in winter barley": S1 sequences

### *EIF4E* wild-type coding sequence, protein sequence and target motifs

M A E D T E T R P A S A G A E E R E E G E I A
ATGGCGGAGGACACGGAGACGAGGCCCGCGTCGGCGGGCGCGGAGGAGAGGGAGGAGGGGGAGATCGCG

D D G D G S A A A A A G R V S A H P L E N A W
GACGACGGAGACGGGTCCGCGGCGGCGGCGGCCGGGCGCGTCAGCGCCCACCCTCTGGAGAACGCCTGG

Target motif CDA1
T F W F D N P Q G K **S** **R** A V A W G S T I H P I
ACCTTCTGGTTCGACAACCCGCAGGGCAAGT**CCC**GGGCGGTGGCCTGGGGGAGCACCATCCACCCCATC

H T F S T V E D F W S L Y N N I H H P S K L N
CACACCTTCTCCACCGTCGAGGACTTCTGGAGCCTTTACAACAATATTCATCACCCTAGCAAGTTGAAT

V G A D F H C F K D K I E P K W E D P I C A N
GTTGGAGCCGACTTCCATTGCTTCAAGGATAAGATTGAGCCAAAATGGGAAGACCCCATTTGTGCCAAT

Target motif ADA2 CDA3
G G **K** W **T** I S C G K G K S D T F W L H T L L A
GGCGGT**AAA**TGGA**CC**ATCAGTTGTGGCAAAGGGAAATCTGACACATTTTGGTTGCATACTTTGCTGGCA

Tar-
L I G E Q F D F G D E I C G A V V S V R K **N** **Q**
TTGATTGGTGAACAATTCGACTTTGGTGATGAAATTTGCGGAGCAGTCGTCAGCGTGCGTAAGAA**CC**AG

get motif CDA4
E R V A I W T K N A A N E T A Q I S I G K Q W
GAAAGAGTAGCTATCTGGACTAAAAATGCTGCCAATGAAACTGCTCAGATAAGCATCGGTAAGCAGTGG

Target motif ADA5
K E F L **D** Y K D S I G F V V H E D A K R S D K
AAGGAGTTTCTGG**A**CTACAAGGACTCCATTGGATTCGTCGTTCATGAGGATGCTAAGAGGTCCGACAAA

G A K N R Y T V *
GGCGCCAAGAACCGCTACACGGTTTGA

### Coding sequences of EIF4E alleles:

>rym4
ATGGCGGAGGACACGGAGACGAGGCCCGCGTCGGCGGGCGCGGAGGAGAGGGAGGAGGGGGAGATCGCGGACGAC
GGAGACGGGTCCGCGGCGGCGGCGGCCGGGCGCGTCAGCGCCCACCCTCTGGAGAACGCCTGGACCTTCTGGTTC
GACAACCCGCAGGGCAAGTTCCGGGCGGTGGCCTGGGGGAGCACCATCCACCCCATCCACACCTTCTCCACCGTC
GAGGACTTCTGGAGCCTTTACAACAATATTCATCACCCTAGCAAGTTGAATGTTGGAGCCGACTTCCATTGCTTC
AAGGATAAGATTGAGCCAAAATGGGAAGACCCCATTTGTGCCAATGGCGGTACATGGACCATCAGTTGTGGCAAA
GGGAAATCTGACACATTTTGGTTGCATACTTTGCTGGCATTGATTGGTGAACAATTCGACTTTGGTGATGAAATT
TGCGGAGCAGTCGTCAGCGTGCGTAAGAACCAGGAAAGAGTAGCTATCTGGACTAAAAATGCTGCCAATGAAACT
GCTCAGATAAGCATCGGTAAGCAGTGGAAGGAGTTTCTGGACTACAAGGACTCCATTGGATTCGTCGTTCATGAG
GATGCTAAGAGGTTCGGCAAAGCCGCCAAGAACCGCTACACGGTTTGA

>rym5
ATGGCGGAGGACACGGAGACGAGGCCCGCGTCGGCGGGCGCGGAGGAGAGGGAGGAGGGGGAGATCGCGGACGAC
GGAGACGGGTCCGCGGCGGCGGCGGCCGGGCGCGTCAGCGCCCACCCTCTGGAGAACGCCTGGACCTTCTGGTTC
GACAACCCGCAGGGCAAGTCCCGGGCGGTGGCCTGGGGGAGCACCATCCACCCCATCCACACCTTCTCCACCGTC
GAGGACTTCTGGAGCCTTTACAACAATATTCATCACCCTAGCAAGTTGAATGTTGGAGCCGACTTCCATTGCTTC
AAGGATAAGATTGAGCCAAAATGGGAAGACCCCATTTGTGCCAATGGCGGTAAATGGAGCATCAGTTGTGGCAAA
GGGAAATCTGACACATTTTGGTTGCATACTTTGCTGGCATTGATTGGTGAACAATTCGACTTTGGTGATGAAATT
TGCGGAGCAGTCGTCAGCGTGCGTAAGGACAAGGAAAGAGTAGCTATCTGGACTAAAAATGCTGCCAATGAAACT
GCTCAGATAAGCATCGGTAAGCAGTGGAAGGAGTTTCTGGACTACAAGGACTCCATTGGATTCGTCGTTCATGAG
GATGCTAAGAGGTCCGACAAAGGCGCCAAGAACCGCTACACGGTTTGA

>$rym_{HOR3298}$
ATGGCGGAGGACACGGAGACGAGGCCCGCGTCGGCGGGCGCGGAGGAGAGGGAGGAGGGGGAGATCGCGGACGAC
GGAGACGGGTCCGCGGCGGCGGCGGCCGGGCGCGTCAGCGCCCACCCTCTGGAGAACGCCTGGACCTTCTGGTTC

GACAACCCGCAGGGCAAGTCCCGGGCGGTGGCCTGGGGGAGCACCATCCACCCCATCCACACCTTCTCCACCGTC
GAGGACTTCTGGAGCCTTTACAACAATATTCATCACCCTAGCAAGTTGAATGTTGGAGCCGACTTCCATTGCTTC
AAGGATAAGATTGAGCCAAAATGGGAAGACCCCATTTGTGCCAATGGCGGTAAATGGACCATCAGTTGTGGCAAA
GGGAAATCTGACACATTTTGGTTGCATACTTTGCTGGCATTGATTGGTGAACAATTCGACTTTGGTGATGAAATT
TGCGGAGCAGTCGTCAGCGTGCGTAAGAACCAGGAAAGAGTAGCTATCTGGACTAAAAATGCTGCCAATGAAACT
GCTCAGATAAGCATCGGTAAGCAGTGGAAGGAGTTTCTGAACTACAAGGACTCCATTGGATTCGTCGTTCATGAG
GATGCTAAGAGGTCCGACAAAGCCGCCAAGAACCGCTACACGGTTTGA

>rymHOR4224
ATGGCGGAGGACACGGAGACGAGGCCCGCGTCGGCGGGCGCGGAGGAGAGGGAGGAGGGGGAGATCGCGGACGAC
GGAGACGGGTCCGCGGCGGCGGCGGCCGGGCGCGTCAGCGCCCACCCTCTGGAGAACGCCTGGACCTTCTGGTTC
GACAACCCGCAGGGCAAGTTCCGGGCGGTGGCCTGGGGGAGCACCATCCACCCCATCCACACCTTCTCCACCGTC
GAGGACTTCTGGAGCCTTTACAACAATATTCATCACCCTAGCAAGTTGAATGTTGGAGCCGACTTCCATTGCTTC
AAGGATAAGATTGAGCCAAAATGGGAAGACCCCATTTGTGCCAATGGCGGTATATGGACCATCAGTTGTGGCAAA
GGGAAATCTGACACATTTTGGTTGCATACTTTGCTGGCATTGATTGGTGAACAATTCGACTTTGGTGATGAAATT
TGCGGAGCAGTCGTCAGCGTGCGTAAGAACCAGGAAAGAGTAGCTATCTGGACTAAAAATGCTGCCAATGAAACT
GCTCAGATAAGCATCGGTAAGCAGTGGAAGGAGTTTCTGGACTACAAGGACTCCATTGGATTCGTCGTTCATGAG
GATGCTAAGAGGTCCGGCAAAGCCGCCAAGAACCGCTACACGGTTTGA

>BE allele 1
ATGGCGGAGGACACGGAGACGAGGCCCGCGTCGGCGGGCGCGGAGGAGAGGGAGGAGGGGGAGATCGCGGACGAC
GGAGACGGGTCCGCGGCGGCGGCGGCCGGGCGCGTCAGCGCCCACCCTCTGGAGAACGCCTGGACCTTCTGGTTC
GACAACCCGCAGGGCAAGTGCCGGGCGGTGGCCTGGGGGAGCACCATCCACCCCATCCACACCTTCTCCACCGTC
GAGGACTTCTGGAGCCTTTACAACAATATTCATCACCCTAGCAAGTTGAATGTTGGAGCCGACTTCCATTGCTTC
AAGGATAAGATTGAGCCAAAATGGGAAGACCCCATTTGTGCCAATGGCGGTAAATGGACCATCAGTTGTGGCAAA
GGGAAATCTGACACATTTTGGTTGCATACTTTGCTGGCATTGATTGGTGAACAATTCGACTTTGGTGATGAAATT
TGCGGAGCAGTCGTCAGCGTGCGTAAGAACCAGGAAAGAGTAGCTATCTGGACTAAAAATGCTGCCAATGAAACT
GCTCAGATAAGCATCGGTAAGCAGTGGAAGGAGTTTCTGGACTACAAGGACTCCATTGGATTCGTCGTTCATGAG
GATGCTAAGAGGTCCGACAAAGCCGCCAAGAACCGCTACACGGTTTGA

>BE allele 2
ATGGCGGAGGACACGGAGACGAGGCCCGCGTCGGCGGGCGCGGAGGAGAGGGAGGAGGGGGAGATCGCGGACGAC
GGAGACGGGTCCGCGGCGGCGGCGGCCGGGCGCGTCAGCGCCCACCCTCTGGAGAACGCCTGGACCTTCTGGTTC
GACAACCCGCAGGGCAAGTCCTGGGCGGTGGCCTGGGGGAGCACCATCCACCCCATCCACACCTTCTCCACCGTC
GAGGACTTCTGGAGCCTTTACAACAATATTCATCACCCTAGCAAGTTGAATGTTGGAGCCGACTTCCATTGCTTC
AAGGATAAGATTGAGCCAAAATGGGAAGACCCCATTTGTGCCAATGGCGGTAAATGGACCATCAGTTGTGGCAAA
GGGAAATCTGACACATTTTGGTTGCATACTTTGCTGGCATTGATTGGTGAACAATTCGACTTTGGTGATGAAATT
TGCGGAGCAGTCGTCAGCGTGCGTAAGAACCAGGAAAGAGTAGCTATCTGGACTAAAAATGCTGCCAATGAAACT
GCTCAGATAAGCATCGGTAAGCAGTGGAAGGAGTTTCTGGACTACAAGGACTCCATTGGATTCGTCGTTCATGAG
GATGCTAAGAGGTCCGACAAAGCCGCCAAGAACCGCTACACGGTTTGA

>BE allele 3
ATGGCGGAGGACACGGAGACGAGGCCCGCGTCGGCGGGCGCGGAGGAGAGGGAGGAGGGGGAGATCGCGGACGAC
GGAGACGGGTCCGCGGCGGCGGCGGCCGGGCGCGTCAGCGCCCACCCTCTGGAGAACGCCTGGACCTTCTGGTTC
GACAACCCGCAGGGCAAGTGCTGGGCGGTGGCCTGGGGGAGCACCATCCACCCCATCCACACCTTCTCCACCGTC
GAGGACTTCTGGAGCCTTTACAACAATATTCATCACCCTAGCAAGTTGAATGTTGGAGCCGACTTCCATTGCTTC
AAGGATAAGATTGAGCCAAAATGGGAAGACCCCATTTGTGCCAATGGCGGTAAATGGACCATCAGTTGTGGCAAA
GGGAAATCTGACACATTTTGGTTGCATACTTTGCTGGCATTGATTGGTGAACAATTCGACTTTGGTGATGAAATT
TGCGGAGCAGTCGTCAGCGTGCGTAAGAACCAGGAAAGAGTAGCTATCTGGACTAAAAATGCTGCCAATGAAACT
GCTCAGATAAGCATCGGTAAGCAGTGGAAGGAGTTTCTGGACTACAAGGACTCCATTGGATTCGTCGTTCATGAG
GATGCTAAGAGGTCCGACAAAGCCGCCAAGAACCGCTACACGGTTTGA

>BE allele 4
ATGGCGGAGGACACGGAGACGAGGCCCGCGTCGGCGGGCGCGGAGGAGAGGGAGGAGGGGGAGATCGCGGACGAC
GGAGACGGGTCCGCGGCGGCGGCGGCCGGGCGCGTCAGCGCCCACCCTCTGGAGAACGCCTGGACCTTCTGGTTC
GACAACCCGCAGGGCAAGTTCTGGGCGGTGGCCTGGGGGAGCACCATCCACCCCATCCACACCTTCTCCACCGTC
GAGGACTTCTGGAGCCTTTACAACAATATTCATCACCCTAGCAAGTTGAATGTTGGAGCCGACTTCCATTGCTTC
AAGGATAAGATTGAGCCAAAATGGGAAGACCCCATTTGTGCCAATGGCGGTAAATGGACCATCAGTTGTGGCAAA
GGGAAATCTGACACATTTTGGTTGCATACTTTGCTGGCATTGATTGGTGAACAATTCGACTTTGGTGATGAAATT
TGCGGAGCAGTCGTCAGCGTGCGTAAGAACCAGGAAAGAGTAGCTATCTGGACTAAAAATGCTGCCAATGAAACT
GCTCAGATAAGCATCGGTAAGCAGTGGAAGGAGTTTCTGGACTACAAGGACTCCATTGGATTCGTCGTTCATGAG
GATGCTAAGAGGTCCGACAAAGCCGCCAAGAACCGCTACACGGTTTGA

>BE allele 5
ATGGCGGAGGACACGGAGACGAGGCCCGCGTCGGCGGGCGCGGAGGAGAGGGAGGAGGGGGAGATCGCGGACGAC
GGAGACGGGTCCGCGGCGGCGGCGGCCGGGCGCGTCAGCGCCCACCCTCTGGAGAACGCCTGGACCTTCTGGTTC
GACAACCCGCAGGGCAAGTTCCGGGTGGTGGCCTGGGGGAGCACCATCCACCCCATCCACACCTTCTCCACCGTC
GAGGACTTCTGGAGCCTTTACAACAATATTCATCACCCTAGCAAGTTGAATGTTGGAGCCGACTTCCATTGCTTC

AAGGATAAGATTGAGCCAAAATGGGAAGACCCCATTTGTGCCAATGGCGGTAAATGGACCATCAGTTGTGGCAAA
GGGAAATCTGACACATTTTGGTTGCATACTTTGCTGGCATTGATTGGTGAACAATTCGACTTTGGTGATGAAATT
TGCGGAGCAGTCGTCAGCGTGCGTAAGAACCAGGAAAGAGTAGCTATCTGGACTAAAAATGCTGCCAATGAAACT
GCTCAGATAAGCATCGGTAAGCAGTGGAAGGAGTTTCTGGACTACAAGGACTCCATTGGATTCGTCGTTCATGAG
GATGCTAAGAGGTCCGACAAAGCCGCCAAGAACCGCTACACGGTTTGA

>BE allele 6
ATGGCGGAGGACACGGAGACGAGGCCCGCGTCGGCGGGCGCGGAGGAGAGGGAGGAGGGGGAGATCGCGGACGAC
GGAGACGGGTCCGCGGCGGCGGCGGCCGGGCGCGTCAGCGCCCACCCTCTGGAGAACGCCTGGACCTTCTGGTTC
GACAACCCGCAGGGCAAGTTCCGGGCGGTGGTCTGGGGGAGCACCATCCACCCCATCCACACCTTCTCCACCGTC
GAGGACTTCTGGAGCCTTTACAACAATATTCATCACCCTAGCAAGTTGAATGTTGGAGCCGACTTCCATTGCTTC
AAGGATAAGATTGAGCCAAAATGGGAAGACCCCATTTGTGCCAATGGCGGTAAATGGACCATCAGTTGTGGCAAA
GGGAAATCTGACACATTTTGGTTGCATACTTTGCTGGCATTGATTGGTGAACAATTCGACTTTGGTGATGAAATT
TGCGGAGCAGTCGTCAGCGTGCGTAAGAACCAGGAAAGAGTAGCTATCTGGACTAAAAATGCTGCCAATGAAACT
GCTCAGATAAGCATCGGTAAGCAGTGGAAGGAGTTTCTGGACTACAAGGACTCCATTGGATTCGTCGTTCATGAG
GATGCTAAGAGGTCCGACAAAGCCGCCAAGAACCGCTACACGGTTTGA

>BE allele 7
ATGGCGGAGGACACGGAGACGAGGCCCGCGTCGGCGGGCGCGGAGGAGAGGGAGGAGGGGGAGATCGCGGACGAC
GGAGACGGGTCCGCGGCGGCGGCGGCCGGGCGCGTCAGCGCCCACCCTCTGGAGAACGCCTGGACCTTCTGGTTC
GACAACCCGCAGGGCAAGTCCCGGGCGGTGGCCTGGGGGAGCACCATCCACCCCATCCACACCTTCTCCACCGTC
GAGGACTTCTGGAGCCTTTACAACAATATTCATCACCCTAGCAAGTTGAATGTTGGAGCCGACTTCCATTGCTTC
AAGGATAAGATTGAGCCAAAATGGGAAGACCCCATTTGTGCCAATGGCGGTAAATGGACCATCAGTTGTGGCAAA
GGGAAATCTGACACATTTTGGTTGCATACTTTGCTGGCATTGATTGGTGAACAATTCGACTTTGGTGATGAAATT
TGCGGAGCAGTCGTCAGCGTGCGTAAGAACGAGGAAAGAGTAGCTATCTGGACTAAAAATGCTGCCAATGAAACT
GCTCAGATAAGCATCGGTAAGCAGTGGAAGGAGTTTCTGGACTACAAGGACTCCATTGGATTCGTCGTTCATGAG
GATGCTAAGAGGTCCGACAAAGCCGCCAAGAACCGCTACACGGTTTGA

>BE allele 8
ATGGCGGAGGACACGGAGACGAGGCCCGCGTCGGCGGGCGCGGAGGAGAGGGAGGAGGGGGAGATCGCGGACGAC
GGAGACGGGTCCGCGGCGGCGGCGGCCGGGCGCGTCAGCGCCCACCCTCTGGAGAACGCCTGGACCTTCTGGTTC
GACAACCCGCAGGGCAAGTCCCGGGCGGTGGCCTGGGGGAGCACCATCCACCCCATCCACACCTTCTCCACCGTC
GAGGACTTCTGGAGCCTTTACAACAATATTCATCACCCTAGCAAGTTGAATGTTGGAGCCGACTTCCATTGCTTC
AAGGATAAGATTGAGCCAAAATGGGAAGACCCCATTTGTGCCAATGGCGGTAAATGGACCATCAGTTGTGGCAAA
GGGAAATCTGACACATTTTGGTTGCATACTTTGCTGGCATTGATTGGTGAACAATTCGACTTTGGTGATGAAATT
TGCGGAGCAGTCGTCAGCGTGCGTAAGAAGGAGGAAAGAGTAGCTATCTGGACTAAAAATGCTGCCAATGAAACT
GCTCAGATAAGCATCGGTAAGCAGTGGAAGGAGTTTCTGGACTACAAGGACTCCATTGGATTCGTCGTTCATGAG
GATGCTAAGAGGTCCGACAAAGCCGCCAAGAACCGCTACACGGTTTGA

>BE allele 9
ATGGCGGAGGACACGGAGACGAGGCCCGCGTCGGCGGGCGCGGAGGAGAGGGAGGAGGGGGAGATCGCGGACGAC
GGAGACGGGTCCGCGGCGGCGGCGGCCGGGCGCGTCAGCGCCCACCCTCTGGAGAACGCCTGGACCTTCTGGTTC
GACAACCCGCAGGGCAAGTCCCGGGCGGTGGCCTGGGGGAGCACCATCCACCCCATCCACACCTTCTCCACCGTC
GAGGACTTCTGGAGCCTTTACAACAATATTCATCACCCTAGCAAGTTGAATGTTGGAGCCGACTTCCATTGCTTC
AAGGATAAGATTGAGCCAAAATGGGAAGACCCCATTTGTGCCAATGGCGGTAAATGGACCATCAGTTGTGGCAAA
GGGAAATCTGACACATTTTGGTTGCATACTTTGCTGGCATTGATTGGTGAACAATTCGACTTTGGTGATGAAATT
TGCGGAGCAGTCGTCAGCGTGCGTAAGAACTAGGAAAGAGTAGCTATCTGGACTAAAAATGCTGCCAATGAAACT
GCTCAGATAAGCATCGGTAAGCAGTGGAAGGAGTTTCTGGACTACAAGGACTCCATTGGATTCGTCGTTCATGAG
GATGCTAAGAGGTCCGACAAAGCCGCCAAGAACCGCTACACGGTTTGA

>BE allele 10
ATGGCGGAGGACACGGAGACGAGGCCCGCGTCGGCGGGCGCGGAGGAGAGGGAGGAGGGGGAGATCGCGGACGAC
GGAGACGGGTCCGCGGCGGCGGCGGCCGGGCGCGTCAGCGCCCACCCTCTGGAGAACGCCTGGACCTTCTGGTTC
GACAACCCGCAGGGCAAGTCCCGGGCGGTGGCCTGGGGGAGCACCATCCACCCCATCCACACCTTCTCCACCGTC
GAGGACTTCTGGAGCCTTTACAACAATATTCATCACCCTAGCAAGTTGAATGTTGGAGCCGACTTCCATTGCTTC
AAGGATAAGATTGAGCCAAAATGGGAAGACCCCATTTGTGCCAATGGCGGTAAATGGACCATCAGTTGTGGCAAA
GGGAAATCTGACACATTTTGGTTGCATACTTTGCTGGCATTGATTGGTGAACAATTCGACTTTGGTGATGAAATT
TGCGGAGCAGTCGTCAGCGTGCGTAAGAAGTAGGAAAGAGTAGCTATCTGGACTAAAAATGCTGCCAATGAAACT
GCTCAGATAAGCATCGGTAAGCAGTGGAAGGAGTTTCTGGACTACAAGGACTCCATTGGATTCGTCGTTCATGAG
GATGCTAAGAGGTCCGACAAAGCCGCCAAGAACCGCTACACGGTTTGA

>BE allele rym4_I
ATGGCGGAGGACACGGAGACGAGGCCCGCGTCGGCGGGCGCGGAGGAGAGGGAGGAGGGGGAGATCGCGGACGAC
GGAGACGGGTCCGCGGCGGCGGCGGCCGGGCGCGTCAGCGCCCACCCTCTGGAGAACGCCTGGACCTTCTGGTTC
GACAACCCGCAGGGCAAGTTCCGGGCGGTGGCCTGGGGGAGCACCATCCACCCCATCCACACCTTCTCCACCGTC

GAGGACTTCTGGAGCCTTTACAACAATATTCATCACCCTAGCAAGTTGAATGTTGGAGCCGACTTCCATTGCTTC
AAGGATAAGATTGAGCCAAAATGGGAAGACCCCATTTGTGCCAATGGCGGTAAATGGACCATCAGTTGTGGCAAA
GGGAAATCTGACACATTTTGGTTGCATACTTTGCTGGCATTGATTGGTGAACAATTCGACTTTGGTGATGAAATT
TGCGGAGCAGTCGTCAGCGTGCGTAAGAACCAGGAAAGAGTAGCTATCTGGACTAAAAATGCTGCCAATGAAACT
GCTCAGATAAGCATCGGTAAGCAGTGGAAGGAGTTTCTGGACTACAAGGACTCCATTGGATTCGTCGTTCATGAG
GATGCTAAGAGGTCCGACAAAGCCGCCAAGAACCGCTACACGGTTTGA

**Protein sequences of EIF4E alleles:**

>rym4
MAEDTETRPASAGAEEREEGEIADDGDGSAAAAAGRVSAHPLENAWTFWFDNPQGKFRAVAWGSTIHPIHTFSTV
EDFWSLYNNIHHPSKLNVGADFHCFKDKIEPKWEDPICANGGTWTISCGKGKSDTFWLHTLLALIGEQFDFGDEI
CGAVVSVRKNQERVAIWTKNAANETAQISIGKQWKEFLDYKDSIGFVVHEDAKRFGKAAKNRYTV*

>rym5
MAEDTETRPASAGAEEREEGEIADDGDGSAAAAAGRVSAHPLENAWTFWFDNPQGKSRAVAWGSTIHPIHTFSTV
EDFWSLYNNIHHPSKLNVGADFHCFKDKIEPKWEDPICANGGKWSISCGKGKSDTFWLHTLLALIGEQFDFGDEI
CGAVVSVRKDKERVAIWTKNAANETAQISIGKQWKEFLDYKDSIGFVVHEDAKRSDKGAKNRYTV*

>rym$_{HOR3298}$
MAEDTETRPASAGAEEREEGEIADDGDGSAAAAAGRVSAHPLENAWTFWFDNPQGKSRAVAWGSTIHPIHTFSTV
EDFWSLYNNIHHPSKLNVGADFHCFKDKIEPKWEDPICANGGKWTISCGKGKSDTFWLHTLLALIGEQFDFGDEI
CGAVVSVRKNQERVAIWTKNAANETAQISIGKQWKEFLNYKDSIGFVVHEDAKRSDKAAKNRYTV*

>rymHOR$_{4224}$
MAEDTETRPASAGAEEREEGEIADDGDGSAAAAAGRVSAHPLENAWTFWFDNPQGKFRAVAWGSTIHPIHTFSTV
EDFWSLYNNIHHPSKLNVGADFHCFKDKIEPKWEDPICANGGIWTISCGKGKSDTFWLHTLLALIGEQFDFGDEI
CGAVVSVRKNQERVAIWTKNAANETAQISIGKQWKEFLDYKDSIGFVVHEDAKRSGKAAKNRYTV*

>BE allele 1
MAEDTETRPASAGAEEREEGEIADDGDGSAAAAAGRVSAHPLENAWTFWFDNPQGKCRAVAWGSTIHPIHTFSTV
EDFWSLYNNIHHPSKLNVGADFHCFKDKIEPKWEDPICANGGKWTISCGKGKSDTFWLHTLLALIGEQFDFGDEI
CGAVVSVRKNQERVAIWTKNAANETAQISIGKQWKEFLDYKDSIGFVVHEDAKRSDKAAKNRYTV*

>BE allele 2
MAEDTETRPASAGAEEREEGEIADDGDGSAAAAAGRVSAHPLENAWTFWFDNPQGKSWAVAWGSTIHPIHTFSTV
EDFWSLYNNIHHPSKLNVGADFHCFKDKIEPKWEDPICANGGKWTISCGKGKSDTFWLHTLLALIGEQFDFGDEI
CGAVVSVRKNQERVAIWTKNAANETAQISIGKQWKEFLDYKDSIGFVVHEDAKRSDKAAKNRYTV*

>BE allele 3
MAEDTETRPASAGAEEREEGEIADDGDGSAAAAAGRVSAHPLENAWTFWFDNPQGKCWAVAWGSTIHPIHTFSTV
EDFWSLYNNIHHPSKLNVGADFHCFKDKIEPKWEDPICANGGKWTISCGKGKSDTFWLHTLLALIGEQFDFGDEI
CGAVVSVRKNQERVAIWTKNAANETAQISIGKQWKEFLDYKDSIGFVVHEDAKRSDKAAKNRYTV*

>BE allele 4
MAEDTETRPASAGAEEREEGEIADDGDGSAAAAAGRVSAHPLENAWTFWFDNPQGKFWAVAWGSTIHPIHTFSTV
EDFWSLYNNIHHPSKLNVGADFHCFKDKIEPKWEDPICANGGKWTISCGKGKSDTFWLHTLLALIGEQFDFGDEI
CGAVVSVRKNQERVAIWTKNAANETAQISIGKQWKEFLDYKDSIGFVVHEDAKRSDKAAKNRYTV*

>BE allele 5
MAEDTETRPASAGAEEREEGEIADDGDGSAAAAAGRVSAHPLENAWTFWFDNPQGKFRVVAWGSTIHPIHTFSTV
EDFWSLYNNIHHPSKLNVGADFHCFKDKIEPKWEDPICANGGKWTISCGKGKSDTFWLHTLLALIGEQFDFGDEI
CGAVVSVRKNQERVAIWTKNAANETAQISIGKQWKEFLDYKDSIGFVVHEDAKRSDKAAKNRYTV*

>BE allele 6
MAEDTETRPASAGAEEREEGEIADDGDGSAAAAAGRVSAHPLENAWTFWFDNPQGKFRAVVWGSTIHPIHTFSTV
EDFWSLYNNIHHPSKLNVGADFHCFKDKIEPKWEDPICANGGKWTISCGKGKSDTFWLHTLLALIGEQFDFGDEI
CGAVVSVRKNQERVAIWTKNAANETAQISIGKQWKEFLDYKDSIGFVVHEDAKRSDKAAKNRYTV*

>BE allele 7
MAEDTETRPASAGAEEREEGEIADDGDGSAAAAAGRVSAHPLENAWTFWFDNPQGKSRAVAWGSTIHPIHTFSTV
EDFWSLYNNIHHPSKLNVGADFHCFKDKIEPKWEDPICANGGKWTISCGKGKSDTFWLHTLLALIGEQFDFGDEI
CGAVVSVRKNEERVAIWTKNAANETAQISIGKQWKEFLDYKDSIGFVVHEDAKRSDKAAKNRYTV*

>BE allele 8
MAEDTETRPASAGAEEREEGEIADDGDGSAAAAAGRVSAHPLENAWTFWFDNPQGKSRAVAWGSTIHPIHTFSTV
EDFWSLYNNIHHPSKLNVGADFHCFKDKIEPKWEDPICANGGKWTISCGKGKSDTFWLHTLLALIGEQFDFGDEI
CGAVVSVRKKEERVAIWTKNAANETAQISIGKQWKEFLDYKDSIGFVVHEDAKRSDKAAKNRYTV*

>BE allele 9
MAEDTETRPASAGAEEREEGEIADDGDGSAAAAAGRVSAHPLENAWTFWFDNPQGKSRAVAWGSTIHPIHTFSTV
EDFWSLYNNIHHPSKLNVGADFHCFKDKIEPKWEDPICANGGKWTISCGKGKSDTFWLHTLLALIGEQFDFGDEI
CGAVVSVRKN*

>BE allele 10
MAEDTETRPASAGAEEREEGEIADDGDGSAAAAAGRVSAHPLENAWTFWFDNPQGKSRAVAWGSTIHPIHTFSTV
EDFWSLYNNIHHPSKLNVGADFHCFKDKIEPKWEDPICANGGKWTISCGKGKSDTFWLHTLLALIGEQFDFGDEI
CGAVVSVRKK*

>BE allele rym4_I
MAEDTETRPASAGAEEREEGEIADDGDGSAAAAAGRVSAHPLENAWTFWFDNPQGKFRAVAWGSTIHPIHTFSTV
EDFWSLYNNIHHPSKLNVGADFHCFKDKIEPKWEDPICANGGKWTISCGKGKSDTFWLHTLLALIGEQFDFGDEI
CGAVVSVRKNQERVAIWTKNAANETAQISIGKQWKEFLDYKDSIGFVVHEDAKRSDKAAKNRYTV*
