## Supplementary material for "Base editing of *EIF4E* creates novel resistance alleles against bymoviruses in winter barley": S2 R code and output

```
### statistical analysis of all homozygous base edited plants ###

### install packages ###

install.packages("tidyverse")
install.packages("AICcmodavg")
install.packages('rstatix')
install.packages('onewaytests')

###open libraries##

library(ggplot2)
library(tidyverse)
library(broom)
library(AICcmodavg)
library(dplyr)
library(onewaytests)
library(rstatix)
```

```
## load data ##
> plant.data <- read.csv("/…/…/260709_EIF4E_Base-Editing_all_data_homozygous.csv", header = TRUE, colClasses = c("fa
ctor", "factor", "factor", "numeric", "numeric", "numeric"))
> summary(plant.data)
                  plant           phenotype          genotype   grain_number   total_grain_weight      TGW
 BPR_42_3_P4-3-2_1 :  1   resistant  :78   rym4_I      :39   Min.   :  0.0   Min.   : 0.000     Min.   :26.50
 BPR_42_3_P4-3-2_10:  1   susceptible:90   wild-type   :31   1st Qu.:182.5   1st Qu.: 8.975     1st Qu.:45.50
 BPR_42_3_P4-3-2_2 :  1                    BE_allele_7:27   Median :240.0   Median :11.300     Median :48.80
 BPR_42_3_P4-3-2_3 :  1                    BE_allele_8:20   Mean   :250.4   Mean   :12.350     Mean   :48.49
 BPR_42_3_P4-3-2_4 :  1                    BE_allele_6:14   3rd Qu.:321.0   3rd Qu.:15.820     3rd Qu.:51.65
 BPR_42_3_P4-3-2_5 :  1                    BE_allele_4:13   Max.   :518.0   Max.   :25.710     Max.   :70.50
 (Other)           :162                    (Other)    :24                                       NAs    :1

> ### A) resistance/ susceptibility ###
>
> ## build cross table ##
>

> crosstable <- table (plant.data$phenotype, plant.data$genotype)
> print(crosstable)
```

```
             BE_allele_1 BE_allele_3 BE_allele_4 BE_allele_6 BE_allele_7 BE_allele_8 BE_allele_9 rym4_I wild-type
  resistant            0           1           8           8          23          16          10      4         8
  susceptible         10           1           5           6           4           4           2     35        23

> ## percentages of resistance ##
>
> per_resistant <- prop.table(crosstable, margin = 2) * 100
> print(round(per_resistant, 2))

             BE_allele_1 BE_allele_3 BE_allele_4 BE_allele_6 BE_allele_7 BE_allele_8 BE_allele_9 rym4_I wild-type
  resistant         0.00       50.00       61.54       57.14       85.19       80.00       83.33  10.26     25.81
  susceptible     100.00       50.00       38.46       42.86       14.81       20.00       16.67  89.74     74.19


>  ## Fisher's Exact Test (qualitative data, variation in sample size between genotypes) ##
>
>   # contigency matrix
>
> contingency_matrix <- table(plant.data$genotype, plant.data$phenotype)
> print(contingency_matrix)

             resistant susceptible
  BE_allele_1         0          10
  BE_allele_3         1           1
  BE_allele_4         8           5
  BE_allele_6         8           6
  BE_allele_7        23           4
  BE_allele_8        16           4
  BE_allele_9        10           2
  rym4_I              4          35
  wild-type           8          23


>   # global Fisher's Exact test
> fisher_result <- fisher.test(contingency_matrix, simulate.p.value = TRUE)
> print(fisher_result)
```

```
        Fisher's Exact Test for Count Data with simulated p-value (based on 2000 replicates)

data:  contingency_matrix
p-value = 0.0004998
alternative hypothesis: two.sided

>   # Post Hoc pairwise Fisher's Exact Test (manual to make it technically more robust)
> 
>    # 1. generate all pairs of genotypes from contigency matrix
> genotypes <- rownames(contingency_matrix)
> pairs <- combn(genotypes, 2)

>    # 2. function to calculate pairwise Fisher test
> run_fisher <- function(g1, g2) {
+   sub_tab <- contingency_matrix[c(g1, g2), ]
+   res <- tryCatch({
+     fisher.test(sub_tab)
+   }, error = function(e) return(NULL))
+ 
+   if(is.null(res)) {
+     return(data.frame(group1 = g1, group2 = g2, p = NA))
+   } else {
+     return(data.frame(group1 = g1, group2 = g2, p = res$p.value))
+   }
+ }

>    # 3. apply function to all pairs of genotypes
> results_list <- apply(pairs, 2, function(x) run_fisher(x[1], x[2]))
> 
>    # 4. convert list into plain data frame
> pairwise_res_manual <- do.call(rbind, results_list)

>    # 5. adjustment of p values according to Benjamin-Hochberg
> valid_idx <- !is.na(pairwise_res_manual$p)
> pairwise_res_manual$p.adj <- NA
> pairwise_res_manual$p.adj[valid_idx] <- p.adjust(pairwise_res_manual$p[valid_idx], method = "BH")
```

```
>    # 6. print results
> print(pairwise_res_manual)
        group1      group2            p        p.adj
1  BE_allele_1 BE_allele_3 1.666667e-01 3.000000e-01
2  BE_allele_1 BE_allele_4 2.716626e-03 8.149879e-03
3  BE_allele_1 BE_allele_6 6.428534e-03 1.780209e-02
4  BE_allele_1 BE_allele_7 2.873711e-06 3.448453e-05
5  BE_allele_1 BE_allele_8 3.331667e-05 1.999000e-04
6  BE_allele_1 BE_allele_9 1.036116e-04 5.328594e-04
7  BE_allele_1      rym4_I 5.686628e-01 6.823954e-01
8  BE_allele_1   wild-type 1.646344e-01 3.000000e-01
9  BE_allele_3 BE_allele_4 1.000000e+00 1.000000e+00
10 BE_allele_3 BE_allele_6 1.000000e+00 1.000000e+00
11 BE_allele_3 BE_allele_7 3.201970e-01 4.802956e-01
12 BE_allele_3 BE_allele_8 4.112554e-01 5.466073e-01
13 BE_allele_3 BE_allele_9 3.956044e-01 5.466073e-01
14 BE_allele_3      rym4_I 2.317073e-01 3.791574e-01
15 BE_allele_3   wild-type 4.772727e-01 5.924765e-01
16 BE_allele_4 BE_allele_6 1.000000e+00 1.000000e+00
17 BE_allele_4 BE_allele_7 1.204319e-01 2.408637e-01
18 BE_allele_4 BE_allele_8 4.251390e-01 5.466073e-01
19 BE_allele_4 BE_allele_9 3.782609e-01 5.446957e-01
20 BE_allele_4      rym4_I 5.456281e-04 2.182512e-03
21 BE_allele_4   wild-type 3.951081e-02 1.015992e-01
22 BE_allele_6 BE_allele_7 6.447835e-02 1.450763e-01
23 BE_allele_6 BE_allele_8 2.522196e-01 3.947786e-01
24 BE_allele_6 BE_allele_9 2.163844e-01 3.709448e-01
25 BE_allele_6      rym4_I 9.972597e-04 3.590135e-03
26 BE_allele_6   wild-type 5.298909e-02 1.271738e-01
27 BE_allele_7 BE_allele_8 7.073302e-01 8.214158e-01
28 BE_allele_7 BE_allele_9 1.000000e+00 1.000000e+00
29 BE_allele_7      rym4_I 6.000094e-10 2.160034e-08
30 BE_allele_7   wild-type 6.120768e-06 4.406953e-05
31 BE_allele_8 BE_allele_9 1.000000e+00 1.000000e+00
32 BE_allele_8      rym4_I 1.463793e-07 2.634827e-06
33 BE_allele_8   wild-type 1.975363e-04 8.889132e-04
34 BE_allele_9      rym4_I 4.284790e-06 3.856311e-05
```

```
35 BE_allele_9   wild-type 1.247947e-03 4.084192e-03
36       rym4_I   wild-type 1.147579e-01 2.408637e-01

> ## stacked plot ##
> plant.data$phenotype <- factor(plant.data$phenotype,
+                                levels = c("susceptible", "resistant"))
> ggplot(plant.data, aes(x = genotype, fill = phenotype)) +
+   geom_bar(position = "stack", width = 0.5) +
+   labs(y = "No. of plants", x = "Genotype", fill = "Phenotype") +
+   theme_bw() +
+   theme(axis.text.x = element_text(angle = 45, hjust = 1)) +
+   scale_fill_manual(values = c("resistant" = "grey35", "susceptible" = "grey90"))
```

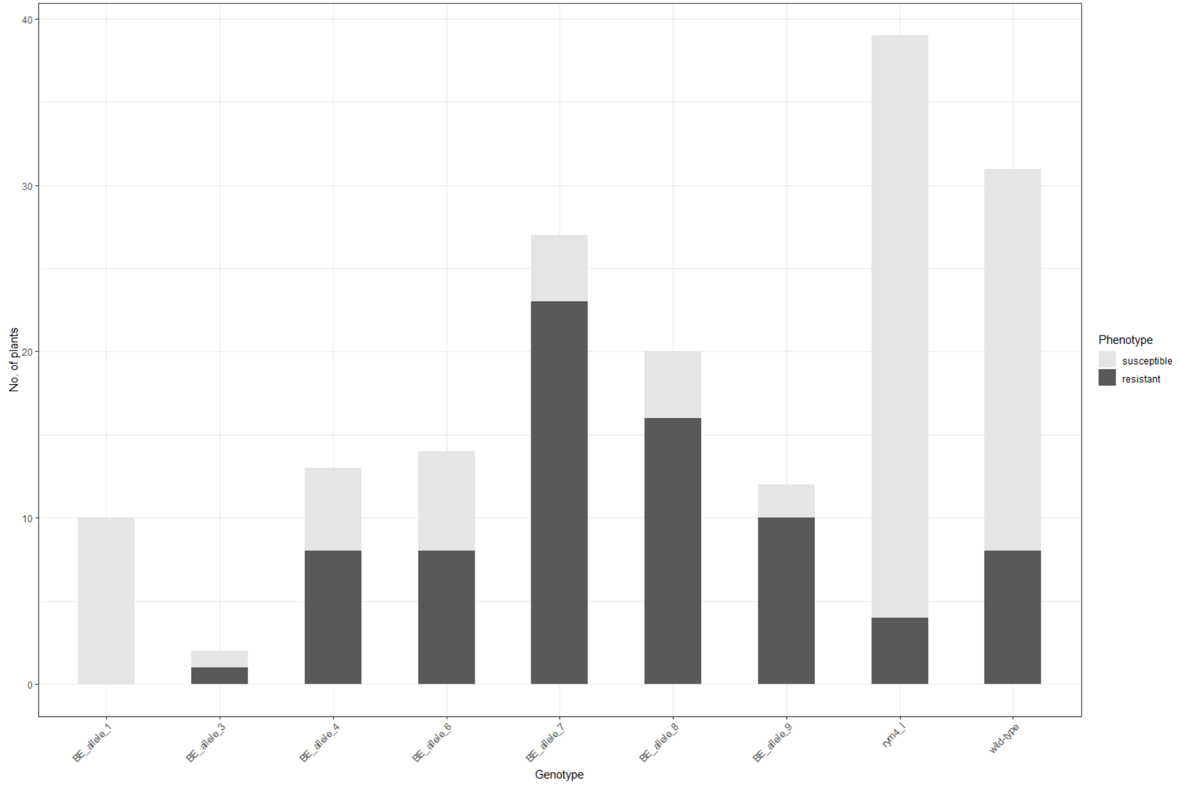

```
> ### B) grain number per plant ###
>
>  ## Test for normal distribution ###
>   # Shapiro-Wilk-Test #
>
> shapiro.test(plant.data$grain_number)

	Shapiro-Wilk normality test

data:  plant.data$grain_number
W = 0.99401, p-value = 0.7262

>   # graphical test with Q-Q-Plot #
>
> qqnorm(plant.data$grain_number)
> qqline(plant.data$grain_number)


> ### Test for equal variance ###
> # Brown-Forsythe-Test #
>
> plant.data %>%
+   group_by(genotype) %>%
+   summarize(var=var(grain_number))
# A tibble: 9 × 2
  genotype       var
  <fct>        <dbl>
1 BE_allele_1  3864.
2 BE_allele_3  5408
3 BE_allele_4  8579.
4 BE_allele_6  3496.
5 BE_allele_7  6628.
6 BE_allele_8  3909.
7 BE_allele_9 12605.
8 rym4_I       4689.
9 wild-type    4911.
>
```

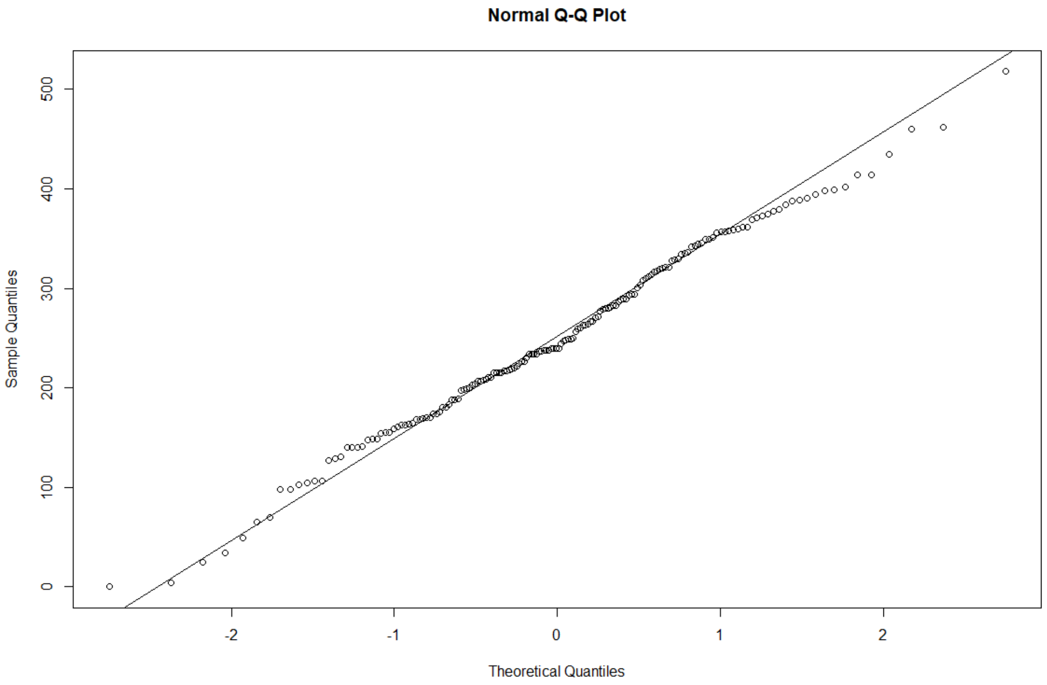

```
> bf.test(grain_number ~ genotype, data = plant.data)

        Brown-Forsythe Test

data:  grain_number and genotype
F = 13.331, num df = 8.0, denom df = 45.8, p-value = 1.019e-09

>  ## Welch ANOVA ## -> Data equally distributed, but high differences in variance
>
> welch_result <- oneway.test(grain_number ~ genotype, data = plant.data)
> print(welch_result)

        One-way analysis of means (not assuming equal variances)

data:  grain_number and genotype
F = 11.802, num df = 8.000, denom df = 19.081, p-value = 6.348e-06


>   # Games-Howell Post-Hoc Test
> gh_ph <- games_howell_test(grain_number ~ genotype, data = plant.data)
Warnmeldung:
There were 2 warnings in `mutate()`.
The first warning was:
i In argument: `data = map(.data$data, .f, ...)`.
Caused by warning in `stats::ptukey()`:
! NaNs wurden erzeugt
 i Run dplyr::last_dplyr_warnings() to see the 1 remaining warning. > print(gh_ph, n= Inf)
# A tibble: 36 × 8
   .y.          group1      group2      estimate   conf.low conf.high       p.adj p.adj.signif
 * <chr>        <chr>       <chr>          <dbl>      <dbl>     <dbl>       <dbl> <chr>
 1 grain_number BE_allele_1 BE_allele_3    -1.30  NaN           NaN    NaN        ""
 2 grain_number BE_allele_1 BE_allele_4   -39.4  -151.           72.2    0.943    "ns"
 3 grain_number BE_allele_1 BE_allele_6    87.8    -0.00709     176.     0.0500   "ns"
 4 grain_number BE_allele_1 BE_allele_7    27.8   -58.7         114.     0.967    "ns"
```

```
 5 grain_number BE_allele_1 BE_allele_8    58.1   -26.3          143.    0.335      "ns"
 6 grain_number BE_allele_1 BE_allele_9  -150.   -283.           -17.1   0.0205     "*"
 7 grain_number BE_allele_1 rym4_I        -50.0  -131.            30.7   0.439      "ns"
 8 grain_number BE_allele_1 wild-type     -60.8  -143.            21.6   0.254      "ns"
 9 grain_number BE_allele_3 BE_allele_4   -38.1   NaN            NaN   NaN          ""
10 grain_number BE_allele_3 BE_allele_6    89.1   NaN            NaN   NaN          ""
11 grain_number BE_allele_3 BE_allele_7    29.1   NaN            NaN   NaN          ""
12 grain_number BE_allele_3 BE_allele_8    59.4   NaN            NaN   NaN          ""
13 grain_number BE_allele_3 BE_allele_9  -149.    NaN            NaN   NaN          ""
14 grain_number BE_allele_3 rym4_I        -48.7   NaN            NaN   NaN          ""
15 grain_number BE_allele_3 wild-type     -59.5   NaN            NaN   NaN          ""
16 grain_number BE_allele_4 BE_allele_6   127.     22.9          232.    0.00997    "**"
17 grain_number BE_allele_4 BE_allele_7    67.2   -36.3          171.    0.421      "ns"
18 grain_number BE_allele_4 BE_allele_8    97.5    -4.28         199.    0.0668     "ns"
19 grain_number BE_allele_4 BE_allele_9  -111.   -253.            31.3   0.213      "ns"
20 grain_number BE_allele_4 rym4_I        -10.6  -109.            88.3   1.000      "ns"
21 grain_number BE_allele_4 wild-type     -21.4  -122.            78.8   0.997      "ns"
22 grain_number BE_allele_6 BE_allele_7   -60.0  -134.            13.5   0.186      "ns"
23 grain_number BE_allele_6 BE_allele_8   -29.7  -100.            40.8   0.885      "ns"
24 grain_number BE_allele_6 BE_allele_9  -238.   -366.          -110.    0.000158   "***"
25 grain_number BE_allele_6 rym4_I       -138.   -203.           -73.0   0.00000381 "****"
26 grain_number BE_allele_6 wild-type    -149.   -216.           -81.1   0.00000123 "****"
27 grain_number BE_allele_7 BE_allele_8    30.3   -38.1           98.7   0.875      "ns"
28 grain_number BE_allele_7 BE_allele_9  -178.   -306.           -50.4   0.00330    "**"
29 grain_number BE_allele_7 rym4_I        -77.8  -140.           -15.8   0.00492    "**"
30 grain_number BE_allele_7 wild-type     -88.6  -154.           -23.5   0.00163    "**"
31 grain_number BE_allele_8 BE_allele_9  -208.   -335.           -81.8   0.000695   "***"
32 grain_number BE_allele_8 rym4_I       -108.   -166.           -50.0   0.0000104  "****"
33 grain_number BE_allele_8 wild-type    -119.   -180.           -57.5   0.00000392 "****"
34 grain_number BE_allele_9 rym4_I        100.    -24.4          225.    0.164      "ns"
35 grain_number BE_allele_9 wild-type      89.5   -36.1          215.    0.276      "ns"
36 grain_number rym4_I      wild-type     -10.8   -64.4           42.8   0.999      "ns"
> dplyr::last_dplyr_warnings()
[[1]]
<warning/rlang_warning>
Warning in `mutate()`:
i In argument: `data = map(.data$data, .f, ...)`.
```

```
Caused by warning in `stats::ptukey()`:
! NaNs wurden erzeugt
---
Backtrace:
    ▆
 1. ├─rstatix::games_howell_test(grain_number ~ genotype, data = plant.data)
 2. │ └─data %>% doo(.games_howell_test, formula, conf.level = conf.level)
 3. ├─rstatix::doo(., .games_howell_test, formula, conf.level = conf.level)
 4. │ └─... %>% mutate(data = map(.data$data, .f, ...))
 5. ├─dplyr::mutate(., data = map(.data$data, .f, ...))
 6. └─dplyr:::mutate.data.frame(., data = map(.data$data, .f, ...))

[[2]]
<warning/rlang_warning>
Warning in `mutate()`:
ℹ In argument: `data = map(.data$data, .f, ...)`.
Caused by warning in `stats::qtukey()`:
! NaNs wurden erzeugt
---
Backtrace:
    ▆
 1. ├─rstatix::games_howell_test(grain_number ~ genotype, data = plant.data)
 2. │ └─data %>% doo(.games_howell_test, formula, conf.level = conf.level)
 3. ├─rstatix::doo(., .games_howell_test, formula, conf.level = conf.level)
 4. │ └─... %>% mutate(data = map(.data$data, .f, ...))
 5. ├─dplyr::mutate(., data = map(.data$data, .f, ...))
 6. └─dplyr:::mutate.data.frame(., data = map(.data$data, .f, ...))

NaNs: → due to low sample size (n=2) for BE allele 3

> ## Boxplot grain number ###
>
> ggplot(data = plant.data) +
+   geom_boxplot(mapping = aes(x = genotype, y = grain_number), width = 0.4) +
+   labs(x = "Genotype", y = "Grain Number") +
+   ylim(0, 600) +
+   theme_bw() +
```

```
+   theme(axis.text.x = element_text(angle = 45, hjust = 1))

> ### C) thousand grain weight ###
>
>  ## Test for normal distribution ###
>   # Shapiro-Wilk-Test #
>
> shapiro.test(plant.data$TGW)

        Shapiro-Wilk normality test

data:  plant.data$TGW
W = 0.95596, p-value = 4.153e-05
```

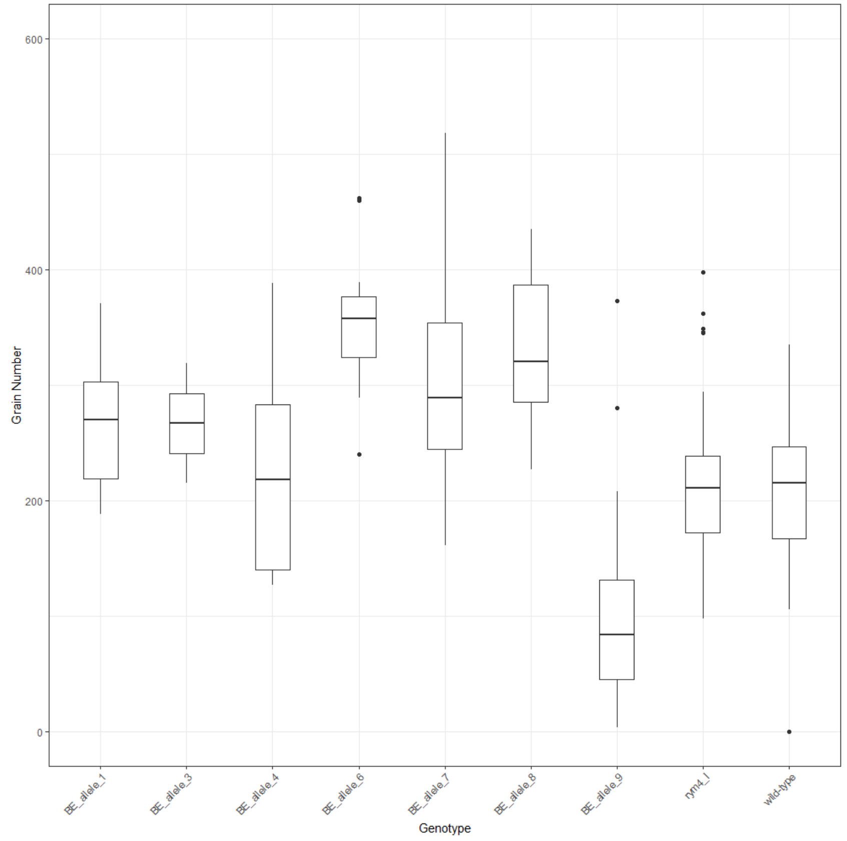

```
>   # graphical test with Q-Q-Plot #
>
> qqnorm(plant.data$TGW)
> qqline(plant.data$TGW)



>  ## Test for normal distribution ###
>   # Shapiro-Wilk-Test #
>
> shapiro.test(plant.data$TGW)

        Shapiro-Wilk normality test

data:  plant.data$TGW
W = 0.82836, p-value = 8.911e-13

> # graphical test with Q-Q-Plot #
>
> qqnorm(plant.data$TGW)
> qqline(plant.data$TGW)


> ### Test for equal variance ###
> # Brown-Forsythe-Test #
>
> plant.data %>%
+   group_by(genotype) %>%
+   summarize(var=var(TGW))
# A tibble: 9 × 2
  genotype      var
  <fct>       <dbl>
1 BE_allele_1  8.33
2 BE_allele_3  7.22
3 BE_allele_4 55.2
4 BE_allele_6 20.2
5 BE_allele_7 15.5
```

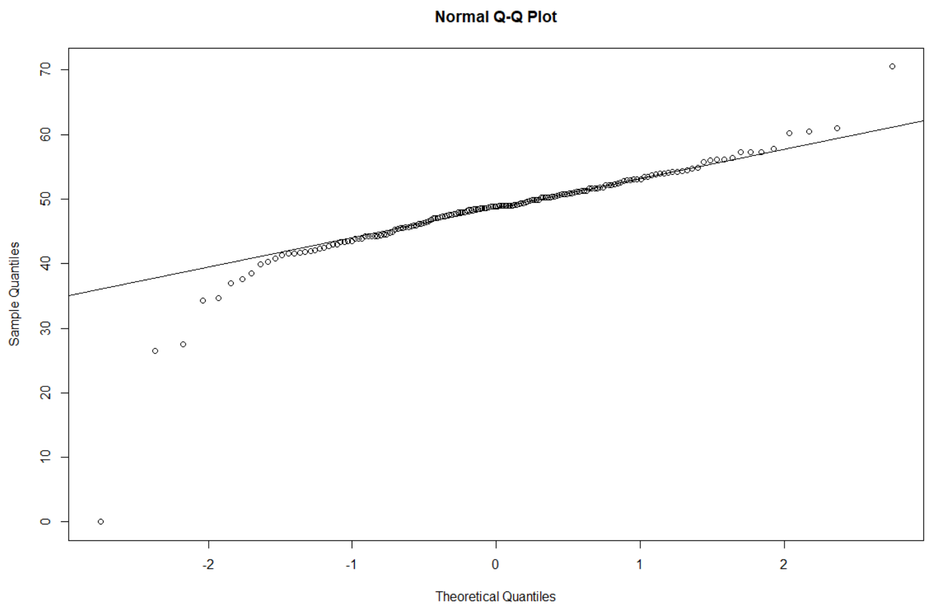

```
6 BE_allele_8 16.6
7 BE_allele_9 48.6
8 rym4_I      16.8
9 wild-type   82.6
>
> bf.test(TGW ~ genotype, data = plant.data)

        Brown-Forsythe Test

data:  TGW and genotype
F = 8.7963, num df = 8.000, denom df = 85.579, p-value = 1.055e-08

>
>
>  ## Kruskal-Wallis-Test ## -> no normal distribution, high differences in variance
>
> # mean and standard distribution of TGW for every genotype
>
> plant.data %>%
+   group_by(genotype) %>%
+   summarise(mean = mean(TGW),
+             sd = sd(TGW))
# A tibble: 9 × 3
  genotype     mean    sd
  <fct>       <dbl> <dbl>
1 BE_allele_1  49.9  2.89
2 BE_allele_3  48.3  2.69
3 BE_allele_4  50.2  7.43
4 BE_allele_6  54.2  4.49
5 BE_allele_7  47.9  3.94
6 BE_allele_8  49.8  4.08
7 BE_allele_9  38.7  6.97
8 rym4_I       49.7  4.10
9 wild-type    45.2  9.09
>
>   #Kruskal-Wallis-Test
> kruskal.test(TGW ~ genotype, data = plant.data)
```

```
        Kruskal-Wallis rank sum test

data:  TGW by genotype
Kruskal-Wallis chi-squared = 49.181, df = 8, p-value = 5.869e-08


>   # Post-hoc-Test for pairwise comparison with p adjustment according to Benjamin-Hochberg
>
> dunn_result <- dunn_test(TGW ~ genotype, data = plant.data,
+                          p.adjust.method = "BH")
> print(dunn_result, n = Inf)
# A tibble: 36 × 9
   .y.   group1      group2         n1    n2 statistic        p       p.adj p.adj.signif
 * <chr> <chr>       <chr>       <int> <int>     <dbl>    <dbl>       <dbl> <chr>
 1 TGW   BE_allele_1 BE_allele_3    10     2   -0.498  6.19e- 1 0.857       ns
 2 TGW   BE_allele_1 BE_allele_4    10    13   -0.306  7.59e- 1 0.911       ns
 3 TGW   BE_allele_1 BE_allele_6    10    14    1.97   4.88e- 2 0.110       ns
 4 TGW   BE_allele_1 BE_allele_7    10    27   -1.08   2.80e- 1 0.439       ns
 5 TGW   BE_allele_1 BE_allele_8    10    20   -0.194  8.46e- 1 0.932       ns
 6 TGW   BE_allele_1 BE_allele_9    10    12   -3.75   1.79e- 4 0.00117     **
 7 TGW   BE_allele_1 rym4_I         10    39   -0.165  8.69e- 1 0.932       ns
 8 TGW   BE_allele_1 wild-type      10    31   -2.08   3.71e- 2 0.0891      ns
 9 TGW   BE_allele_3 BE_allele_4     2    13    0.338  7.35e- 1 0.911       ns
10 TGW   BE_allele_3 BE_allele_6     2    14    1.59   1.12e- 1 0.212       ns
11 TGW   BE_allele_3 BE_allele_7     2    27   -0.0195 9.84e- 1 0.984       ns
12 TGW   BE_allele_3 BE_allele_8     2    20    0.419  6.75e- 1 0.868       ns
13 TGW   BE_allele_3 BE_allele_9     2    12   -1.60   1.10e- 1 0.212       ns
14 TGW   BE_allele_3 rym4_I          2    39    0.451  6.52e- 1 0.868       ns
15 TGW   BE_allele_3 wild-type       2    31   -0.511  6.10e- 1 0.857       ns
16 TGW   BE_allele_4 BE_allele_6    13    14    2.45   1.42e- 2 0.0393      *
17 TGW   BE_allele_4 BE_allele_7    13    27   -0.802  4.22e- 1 0.633       ns
18 TGW   BE_allele_4 BE_allele_8    13    20    0.151  8.80e- 1 0.932       ns
19 TGW   BE_allele_4 BE_allele_9    13    12   -3.69   2.28e- 4 0.00117     **
20 TGW   BE_allele_4 rym4_I         13    39    0.220  8.26e- 1 0.932       ns
21 TGW   BE_allele_4 wild-type      13    31   -1.90   5.69e- 2 0.120       ns
22 TGW   BE_allele_6 BE_allele_7    14    27   -3.69   2.23e- 4 0.00117     **
```

```
23 TGW   BE_allele_6 BE_allele_8     14    20  -2.56    1.06e- 2 0.0346       *
24 TGW   BE_allele_6 BE_allele_9     14    12  -6.15    7.64e-10 0.0000000275 ****
25 TGW   BE_allele_6 rym4_I          14    39  -2.81    5.01e- 3 0.0180       *
26 TGW   BE_allele_6 wild-type       14    31  -4.89    1.02e- 6 0.0000184    ****
27 TGW   BE_allele_7 BE_allele_8     27    20   1.10    2.71e- 1 0.439        ns
28 TGW   BE_allele_7 BE_allele_9     27    12  -3.47    5.16e- 4 0.00232      **
29 TGW   BE_allele_7 rym4_I          27    39   1.36    1.73e- 1 0.297        ns
30 TGW   BE_allele_7 wild-type       27    31  -1.36    1.73e- 1 0.297        ns
31 TGW   BE_allele_8 BE_allele_9     20    12  -4.19    2.81e- 5 0.000253     ***
32 TGW   BE_allele_8 rym4_I          20    39   0.0601  9.52e- 1 0.979        ns
33 TGW   BE_allele_8 wild-type       20    31  -2.38    1.72e- 2 0.0443       *
34 TGW   BE_allele_9 rym4_I          12    39   4.68    2.82e- 6 0.0000339    ****
35 TGW   BE_allele_9 wild-type       12    31   2.49    1.28e- 2 0.0384       *
36 TGW   rym4_I      wild-type       39    31  -2.91    3.65e- 3 0.0146       *

>   # calculation of effect size
> 
> efs <- dunn_test(TGW ~ genotype, data=plant.data,
+                p.adjust.method = "BH")
> efs$r <- efs$statistic/sqrt(d$n1+d$n2)
> print(efs, n = Inf)
# A tibble: 36 × 10
   .y.   group1      group2         n1    n2 statistic        p   p.adj p.adj.signif        r
   <chr> <chr>       <chr>       <int> <int>     <dbl>    <dbl>   <dbl> <chr>           <dbl>
 1 TGW   BE_allele_1 BE_allele_3    10     2  -0.498   6.19e- 1 0.857   ns           -0.144
 2 TGW   BE_allele_1 BE_allele_4    10    13  -0.306   7.59e- 1 0.911   ns           -0.0639
 3 TGW   BE_allele_1 BE_allele_6    10    14   1.97    4.88e- 2 0.110   ns            0.402
 4 TGW   BE_allele_1 BE_allele_7    10    27  -1.08    2.80e- 1 0.439   ns           -0.178
 5 TGW   BE_allele_1 BE_allele_8    10    20  -0.194   8.46e- 1 0.932   ns           -0.0354
 6 TGW   BE_allele_1 BE_allele_9    10    12  -3.75    1.79e- 4 0.00117 **           -0.799
 7 TGW   BE_allele_1 rym4_I         10    39  -0.165   8.69e- 1 0.932   ns           -0.0236
 8 TGW   BE_allele_1 wild-type      10    31  -2.08    3.71e- 2 0.0891  ns           -0.330
 9 TGW   BE_allele_3 BE_allele_4     2    13   0.338   7.35e- 1 0.911   ns            0.0872
10 TGW   BE_allele_3 BE_allele_6     2    14   1.59    1.12e- 1 0.212   ns            0.397
11 TGW   BE_allele_3 BE_allele_7     2    27  -0.0195  9.84e- 1 0.984   ns           -0.00362
12 TGW   BE_allele_3 BE_allele_8     2    20   0.419   6.75e- 1 0.868   ns            0.0892
13 TGW   BE_allele_3 BE_allele_9     2    12  -1.60    1.10e- 1 0.212   ns           -0.427
```

```
14 TGW   BE_allele_3 rym4_I         2    39   0.451  6.52e- 1 0.868         ns          0.0704
15 TGW   BE_allele_3 wild-type      2    31  -0.511  6.10e- 1 0.857         ns         -0.0903
16 TGW   BE_allele_4 BE_allele_6   13    14   2.45   1.42e- 2 0.0393        *           0.472
17 TGW   BE_allele_4 BE_allele_7   13    27  -0.802  4.22e- 1 0.633         ns         -0.127
18 TGW   BE_allele_4 BE_allele_8   13    20   0.151  8.80e- 1 0.932         ns          0.0263
19 TGW   BE_allele_4 BE_allele_9   13    12  -3.69   2.28e- 4 0.00117       **         -0.737
20 TGW   BE_allele_4 rym4_I        13    39   0.220  8.26e- 1 0.932         ns          0.0305
21 TGW   BE_allele_4 wild-type     13    31  -1.90   5.69e- 2 0.120         ns         -0.290
22 TGW   BE_allele_6 BE_allele_7   14    27  -3.69   2.23e- 4 0.00117       **         -0.576
23 TGW   BE_allele_6 BE_allele_8   14    20  -2.56   1.06e- 2 0.0346        *          -0.438
24 TGW   BE_allele_6 BE_allele_9   14    12  -6.15   7.64e-10 0.000000275   ****       -1.21
25 TGW   BE_allele_6 rym4_I        14    39  -2.81   5.01e- 3 0.0180        *          -0.385
26 TGW   BE_allele_6 wild-type     14    31  -4.89   1.02e- 6 0.0000184     ****       -0.737
27 TGW   BE_allele_7 BE_allele_8   27    20   1.10   2.71e- 1 0.439         ns          0.161
28 TGW   BE_allele_7 BE_allele_9   27    12  -3.47   5.16e- 4 0.00232       **         -0.556
29 TGW   BE_allele_7 rym4_I        27    39   1.36   1.73e- 1 0.297         ns          0.168
30 TGW   BE_allele_7 wild-type     27    31  -1.36   1.73e- 1 0.297         ns         -0.180
31 TGW   BE_allele_8 BE_allele_9   20    12  -4.19   2.81e- 5 0.000253      ***        -0.740
32 TGW   BE_allele_8 rym4_I        20    39   0.0601 9.52e- 1 0.979         ns          0.00782
33 TGW   BE_allele_8 wild-type     20    31  -2.38   1.72e- 2 0.0443        *          -0.337
34 TGW   BE_allele_9 rym4_I        12    39   4.68   2.82e- 6 0.0000339     ****        0.656
35 TGW   BE_allele_9 wild-type     12    31   2.49   1.28e- 2 0.0384        *           0.384
36 TGW   rym4_I      wild-type     39    31  -2.91   3.65e- 3 0.0146        *          -0.350

> ggplot(data = plant.data) +
+   geom_boxplot(mapping = aes(x = genotype, y = TGW), width = 0.4) +
+   labs(x = "Genotype", y = "Thousand Grain Weight") +
+   ylim(0, 80) +
+   theme_bw() +
+   theme(axis.text.x = element_text(angle = 45, hjust = 1))
```

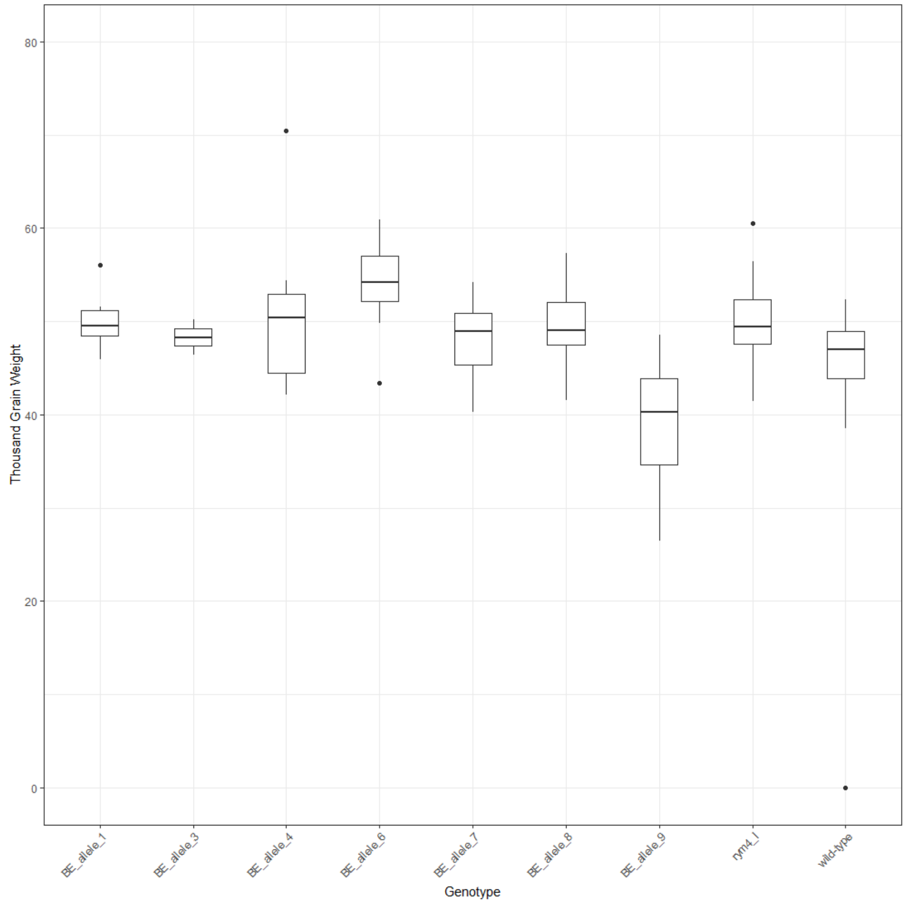
Thousand Grain Weight
80
60
40
20
0
BE_allele_1
BE_allele_3
BE_allele_4
BE_allele_6
BE_allele_7
BE_allele_8
BE_allele_9
rym4_l
wild-type
Genotype

>
